# Oxidative Stress and Interferon Signaling Drive Differential Pathogenesis of Ancestral and Contemporary Zika Viruses in Human Cerebral Organoids

**DOI:** 10.1101/2025.08.12.669648

**Authors:** Alfred T. Harding, Yichen Zhang, Jane-Jane Chen, Jenna M. Antonucci, Alexsia Richards, Valerie Leger, Charles A. Whittaker, Yann S. Vanrobaeys, Divyansh Agarwal, Tenzin Lungjangwa, Rudolf Jaenisch, Lee Gehrke

## Abstract

Neurotropic Zika viruses (ZIKV) cause serious human disease with pandemic potential. Pathogenesis severities resulting from Asian/American versus African ZIKV lineage infections range from mild to severe, respectively; however, mechanisms underlying differential ZIKV pathogenesis remain unclear, as do effective therapeutic strategies. The limitations of mechanistic understanding are due in part to the challenges of comparing data generated in disparate experimental models, as well as approaches that did not test both ancestral and contemporary ZIKV infections. The goal of this work was to define differential pathogenesis mechanisms among ancestral and contemporary ZIKVs by direct infection comparisons using a relevant human stem cell-derived cerebral organoid experimental model. While Asian/American ZIKV lineage infections enhanced antiviral and interferon gene expression responses that correlated with viral RNA clearance from organoid ventricles, ancestral African lineage ZIKV infections enhanced apoptotic and stress response signaling that correlated with diminished STAT2 signaling protein levels, ongoing ZIKV replication, and production of damaging reactive oxygen species (ROS). We discovered that, surprisingly, severe ancestral Zika virus pathogenesis was dramatically reduced by Trolox, a hydroxyl radical scavenger antioxidant, thereby confirming ROS imbalance as a major pathogenesis driver. These results demonstrate that ZIKV lineage infections and pathogenesis are differentiated by their signaling responses and suggest that preventing or controlling hydroxyl radical imbalance may offer therapeutic benefits to address microcephaly and Congenital Zika Virus Syndrome.

**One Sentence Summary:** Differential signal transduction responses to lineage-specific Zika virus infections cause reduction-oxidation imbalance-mediated pathogenesis that is blocked by Trolox, an antioxidant.

## INTRODUCTION

Zika virus (ZIKV) is a mosquito-transmitted positive sense single-stranded RNA virus and a member of the virus family *Flaviviridae*. ZIKV was first isolated from a sentinel monkey in Africa in 1947 (*1*), and two ZIKV lineages (African and Asian) were described in 2012 (*2*), with a third lineage (American) included in 2019 (*3, 4*). Fewer than twenty ZIKV infections were recorded prior to 2007 (*5*), when outbreaks occurred in the Yap Islands and French Polynesia (*6*) with Guillain-Barré syndrome occurrence (*7*). In 2015, an epidemic of infections was reported in the Americas, especially in northeastern Brazil (*8*), along with reports of fetal microcephaly and adult Guillain-Barré syndrome (*6, 9–11*). Zika virus is unusual among flaviviruses in its ability to cross placental barriers (*12–14*), and ZIKV infections have been correlated with an array of congenital abnormalities, including microcephaly and Congenital Zika Syndrome (CZS).

Viral pathogenesis is a complex, multifaceted process that pits viral RNA replication against host antiviral responses. Host cells have effective antiviral responses that degrade cellular and viral RNA (OAS-RNaseL) (*15*), inhibit protein synthesis (eIF2α kinases) (*16*), inhibit the viral RNA-dependent RNA polymerase (MX proteins) (*17*), or block viral RNA translation (IFIT proteins) (*18*). To counter these host responses, viruses can disrupt viral RNA sensing pathways (*19, 20*), block interferon production (*21, 22*), prevent apoptosis (*23*), or disrupt signal transduction (*92*). In the face of a viral infection, host cells activate stress responses that function to maintain homeostasis and facilitate recovery from a virus attack. Stress responses have been correlated with ZIKV infections (*24–26*), but their causative roles across ZIKV with differential pathogenesis are not understood.

Viral protein amino acid substitutions that preceded ZIKV spread from Africa and Asia to the Americas were initially proposed to be causal in the context of enhanced ZIKV transmission, pathogenesis, or microcephaly (*27–29*). A prM protein S139N substitution was correlated with microcephaly in a mouse model (*27*), but recent experimentation has not corroborated the findings (*29, 30*). It has been reported that during spread into the Americas, the Brazilian ZIKV strains acquired nucleotide substitutions that increased viral virulence (*31, 32*); however, others found that African lineages were more pathogenic in both cell culture and murine models (*13, 33-34*). In general, agreement across the field on ZIKV pathogenesis has been limited by attempts to compare or extrapolate use of different model systems, including transformed cell lines, stem cell-derived cells, rodents, and non-human primates, as well as experimentation that did not include ZIKVs from all lineages.

We reasoned that a side-by-side comparison of ZIKV infections, using a relevant human model system, would inform an understanding of ZIKV pathogenesis and disease mechanisms. Cerebral organoids derived from human pluripotent stem cells recapitulate gene expression patterns and brain morphology during fetal development (*35–38*), and have been shown to be an excellent model for studying viral neuropathogenesis (*37, 39, 40*). Here, we have systematically infected human cerebral organoids with a panel of ZIKV lineages and strains collected from several geographic locations, mirroring ZIKV transmission from Africa to Asia and the Americas. Organoid size measurements and confocal immunofluorescence imaging of internal organoid structures were employed to examine ventricles and cytoarchitecture. Bulk and single cell RNA sequencing, protein immunoblotting, and biochemical assays were used to analyze processes underlying ZIKV pathogenesis, which ranged from minor cytoarchitecture disruption to growth arrest with severe disruption of the organoid cytoarchitecture. The results demonstrate that contemporary ZIKV pathogenesis mechanisms include differential positive enrichment of antiviral and interferon gene expression pathways, while severe ZIKV pathogenesis is characterized by positive enrichment of apoptotic and cellular stress gene expression pathways, elevated mitochondrial superoxide levels, and loss of the key STAT2 signaling protein. Of critical significance, severe ZIKV pathogenesis was largely prevented in organoids by Trolox, a vitamin E analog hydroxyl radical scavenger and antioxidant.

## RESULTS

The phylogenetic relationships of the seven ZIKV strains used in this study are summarized in Fig. 1A, showing two Asian lineage strains (Malaysia and Cambodia), two American lineage strains (Puerto Rico and Brazil), and three African lineage strains (Senegal, Nigeria, Uganda). Cerebral organoids at days 1-21 post-embryoid body (EB) formation (Fig. 1B) correlate well with structural and molecular data describing first trimester human brain development (*41*). Organoids were measured following infection with seven ZIKV strains (Fig. 2A) and the data were quantified over a 21-day period (Fig. 2B). The results demonstrate that Asian/American ZIKV strains infections did not significantly affect organoid growth; however, African ZIKV strains (ZIKV-UG, ZIKV-SEN, and ZIKV-NIG) slowed organoid growth, plateauing by 14 dpi (Fig. 2B). Although ZIKV-MAL is classified as an Asian lineage virus, its growth curves also plateaued between days 14 and 21 (Fig. 2B), similar to the African lineage infections. When plotted as change in organoid size per day, the results show that the organoids infected by African lineage ZIKV were smaller at 21 dpi than at 14 dpi (Fig. 2C), suggesting cell death. Others have reported that organoids infected by the African MR766 strain grow more slowly than uninfected human cerebral organoids (*37, 39, 40*). Plaque assays demonstrated that ZIKV-PR-infected organoids released 10-15-fold fewer plaque forming units (PFU) of ZIKV-PR at 7 dpi as compared to all other ZIKV strains (Fig. 2D). By 14 and 21 dpi, the PFU levels from organoids infected by the Asian/American lineage ZIKV (ZIKV-PR, ZIKB-BR, and ZIKV-CAM) decreased to levels approximately 2-logs lower than ZIKV-MAL and the African lineage viruses, remaining constant from 7 dpi. Overall, all ZIKV-infected organoids have decreased growth compared to Mock, indicating slowed growth, but only the African ZIKVs had negative growth. One hypothesis to explain these results is that the Asian/American ZIKV replication and spread were controlled by host anti-viral responses that were less effective in organoids infected by ZIKV-MAL and African lineage viruses, leading to higher levels of virus release.

**Figure 1.**
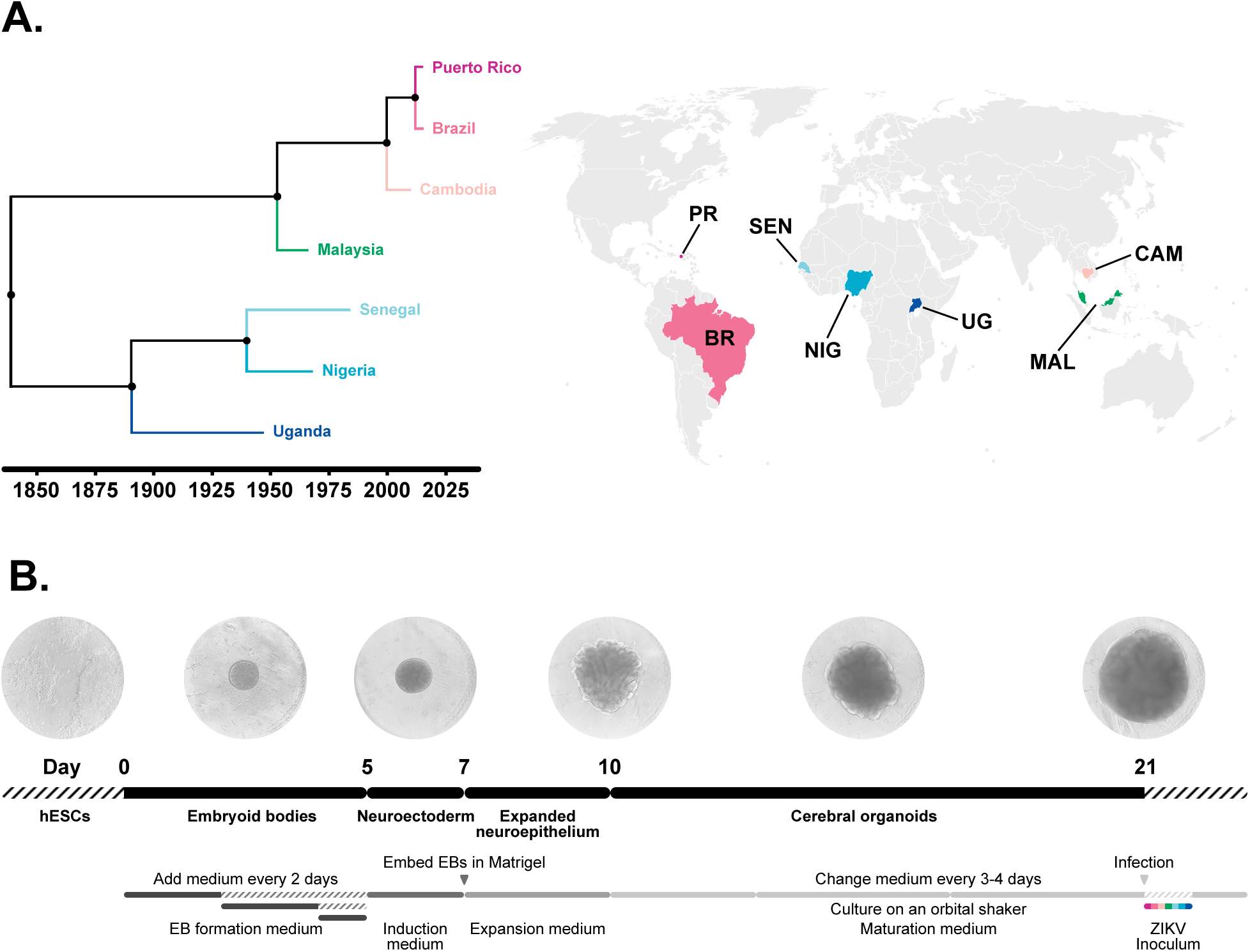
Dendrogram and organoid formation timeline. **A)** Dendrogram of the seven ZIKV strains generated with BEAST (version 2.2.1) demonstrating clustering of the African lineages (UG, NIG, SEN), and the contemporary Asian and American strains (PR, BRA, CAM, MAL). World map showing geographical distribution of the chosen ZIKV panel. **B)** Cerebral organoid formation and infection timeline with representative brightfield microscope images at 3, 6, 10, 14, and 21 days post-EB formation.

**Figure 2.**
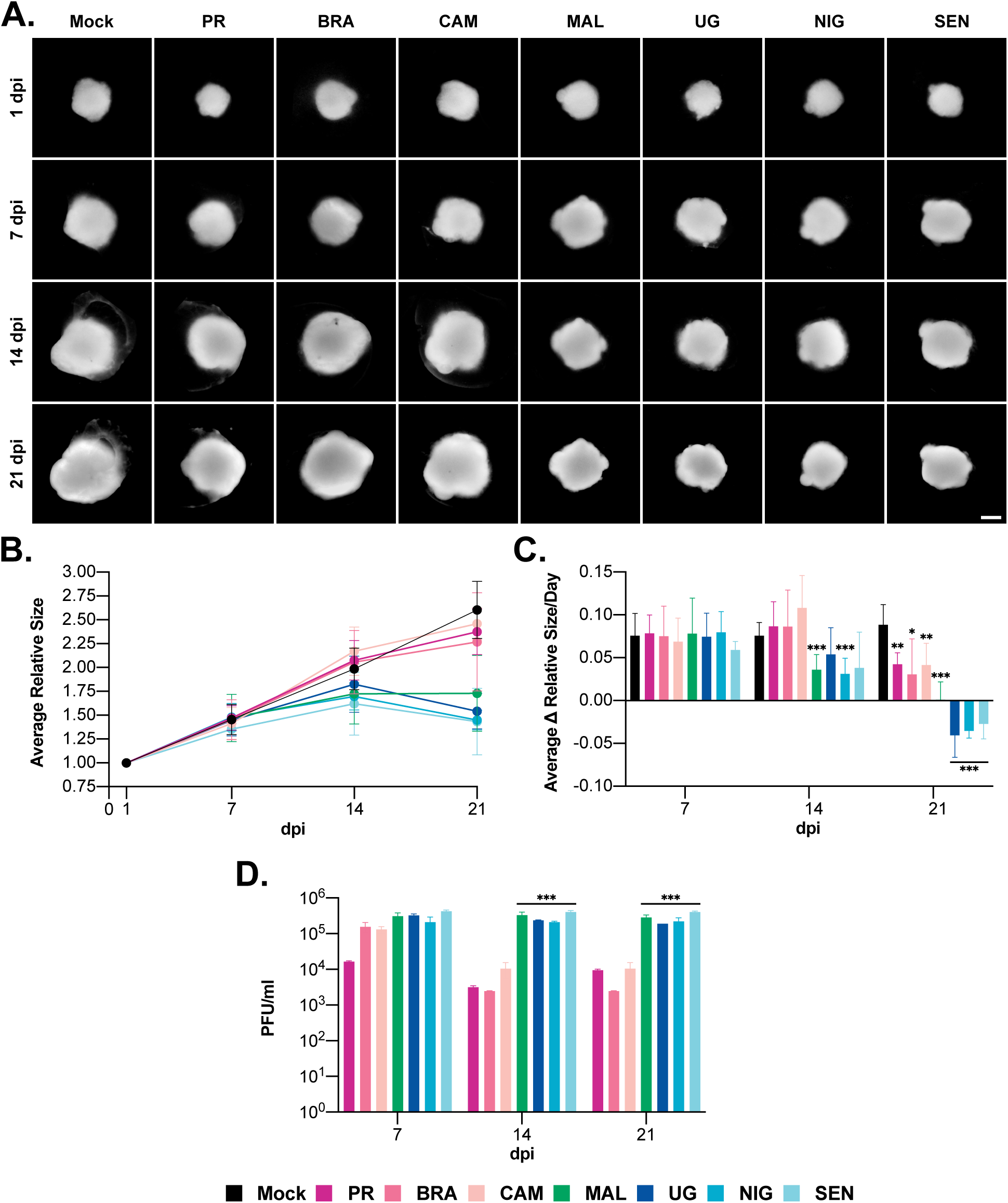
Growth characteristics of cerebral organoids infected with ZIKVs. **A)** Darkfield microscope images of organoids at 1, 7, 14, and 21 dpi (scale bar = 500 μm). **B)** Average size of organoids in each infection group relative to 1 dpi at 7, 14, and 21 dpi (n = 9; mean ± SD). **C)** Average change in size/day at 7, 14, and 21 dpi of organoids in each infection group (n = 9, *p ≤ 0.05, **p < 0.01, ***p < 0.001 compared to Mock; mean ± SD). **D)** Viral titers representing 24 hrs of released virus at 7, 14, and 21 dpi (n = 3, ***p < 0.001 compared to PR, BR, and CAM; mean ± SD).

### ZIKV-induced growth defect severity correlates with cytoarchitectural disruption

We next cryosectioned the organoids and performed immunofluorescence imaging using anti-SRY-box 2 (SOX2; recognizes the SOX2 transcription factor and identifies progenitor cells), anti-microtubule associated protein 2 (MAP2; recognizes the neuronal soma and dendrites), and anti-double stranded RNA (J2; recognizes the double stranded viral replicative intermediate RNA) antibodies. A schematic of an organoid section is presented in Fig. S1, showing that ventricles are composed of neuronal progenitor cells, intermediate progenitor cells, and radial glial; these cells are referred to here as ventricle cells. Glia, astrocytes, and oligodendrocytes are found outside of the ventricles. Organoid ventricles are located near the periphery of organoid sections and are analogous to human brain ventricles. Several progenitor cell types are recognized by anti-SOX2 antibodies, including radial glia, neural progenitor cells (NPC), and intermediate progenitor cells (Fig. S1). SOX2 and MAP2 antibody reactivity is apparent in ventricles of 7 dpi organoids (Fig. 3). The mock-infected human cerebral organoids and the Asian/American strain-infected organoids (except ZIKV-MAL) have similar morphologies, with SOX2+ ventricles surrounded by MAP2+ neurons (*42*) (Fig. S1; Fig. 3). At 14 dpi, the ventricle sizes increased in the mock-infected organoids while the total number of ventricles decreased. These ventricles were maintained through 21 dpi, along with the surrounding interlaced meshwork of MAP2+ neurons. Conversely, at 14 dpi, organoids infected by ZIKV-MAL, ZIKV-UG, ZIKV-NIG and ZIKV-SEN showed cytoarchitectural disorganization and low levels of SOX2 staining, suggesting loss of neuronal progenitor cells (*43*). ZIKV-MAL, ZIKV-UG, ZIKV-NIG and ZIKV-SEN -infected organoids displayed a relative absence of the neuropil (dense meshwork of axons and dendrites) at 14 dpi and 21 dpi. The immunofluorescence data suggest that the organoid growth defects (Fig. 2) correlate with loss of SOX2+ cells and disruption of the MAP2+ organoid neuronal neuropil and cytoarchitecture (Fig. 3).

**Figure 3.**
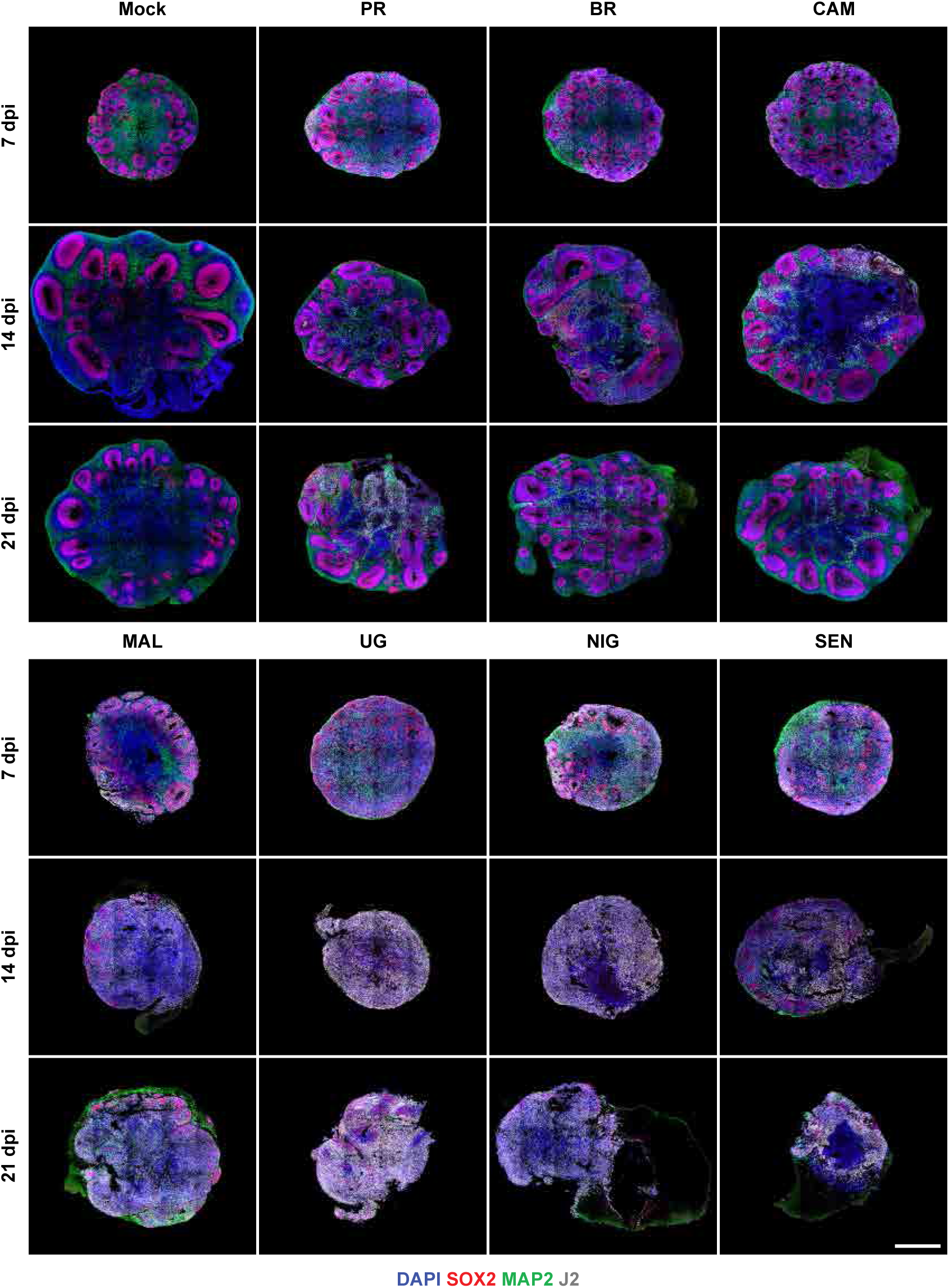
Confocal imaging of cerebral organoids infected by seven ZIKV strains at 7, 14, and 21 dpi. The immunofluorescence signal is from anti-SOX2 antibody (red, progenitor cells), anti-MAP2 antibody (green, neuron somata and dendrites), anti-double stranded viral replicative RNA antibody (white, J2), and DAPI (blue). The ZIKV strain abbreviations are: PR: Puerto Rico; BRA: Brazil; CAM: Cambodia; MAL: Malaysia; UG: Uganda; NIG: Nigeria; SEN: Senegal. The scale bar represents 500 μm.

### ZIKV RNA is cleared from SOX2-positive ventricle cells in Asian/American strain infections, but not in African lineage ZIKV and ZIKV-MAL infections

Organoid sections were viewed at higher magnification to examine ventricle organization and quantify virus-infected SOX2+ ventricle cells (Fig. 4). SOX2 only staining is shown in Fig 4 (rows A, C, E) while the corresponding images including the J2 channel are presented in Fig. rows B, D, F. Quantification of SOX2+/J2+ ventricle cells is shown in Fig. 4G. Seven dpi organoid ventricles infected by ZIKV-PR, ZIKV-BR, or ZIKV-CAM showed 60% infection in SOX2+ cells (Fig. 4 row B, columns 2-4; Fig. 4G). Strikingly, the percentages of ventricular SOX2+/J2+ cells observed in the ZIKV-PR, ZIKV-BR, and ZIKV-CAM -infected organoids decreased precipitously by 14 dpi and remained low through 21 dpi (Fig. 4, rows B, D, and F, columns 2-4; Fig. 4G). J2 signal was also diminished in the extraventricular areas that are expected to be occupied by mature neurons, astrocytes, and oligodendrocytes (Fig. 4, rows D and F, columns 2-4), but not as pronounced as in the ventricles. The J2 signal persisted in ventricular progenitor cells infected by ZIKV-MAL and the African ZIKV strains (ZIKV-UG, ZIKV-NIG, and ZIKV-SEN), remaining at 70-90% over the time course (Fig. 4, rows B, D, and F, columns 5-8; Fig. 4G). These results suggest that replication and spread of Asian/American ZIKV lineage viruses were limited in ventricle cells, while replication and spread were seemingly unimpeded in ventricle cells infected by ZIKV-MAL and the African lineage ZIKV strains.

**Figure 4.**
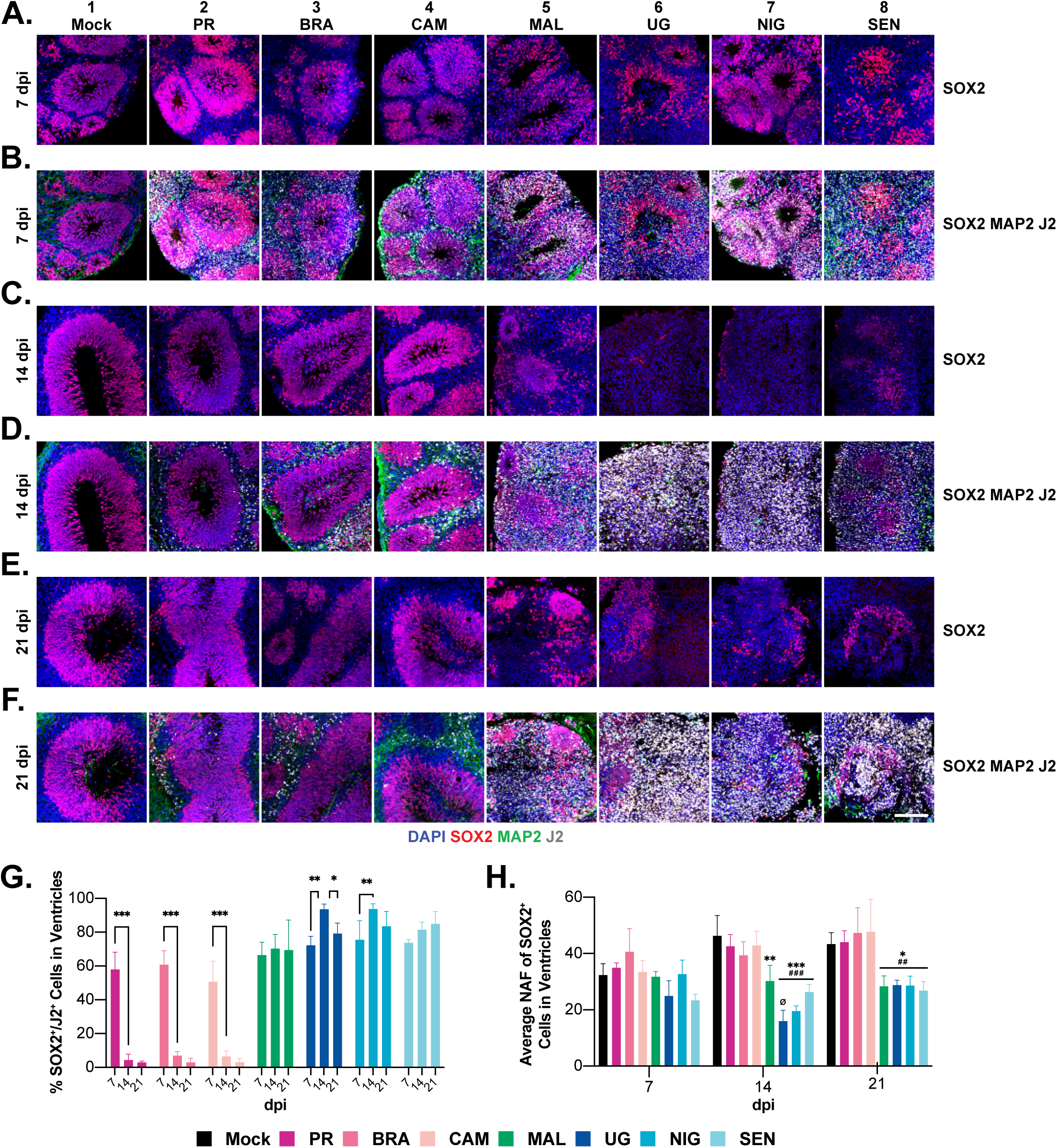
Immunofluorescence microscopy visualizes ZIKV infection persistence in SOX2-positive cells. Confocal microscopy images showing ventricle regions of human cerebral organoids that were mock-infected or infected with ZIKV and then sectioned and stained with anti-SOX2 antibody (red, progenitor cells), anti-MAP2 antibody (green, neuron somata and dendrites), anti-double stranded viral replicative RNA antibody (white, J2), and DAPI (blue) at 7, 14, and 21 dpi (scale bar = 100 μm). The ZIKV strain abbreviations are: PR: Puerto Rico; BRA: Brazil; CAM: Cambodia; MAL: Malaysia; UG: Uganda; NIG: Nigeria; SEN: Senegal. For G-H, the statistical annotation is: n = 3, *p ≤ 0.05, **p < 0.01, ***p < 0.001, * compared to Mock, ^#^ compared to PR, BRA, and CAM, ^Ø^ compared to MAL; mean ± SD. **A)** 7 dpi SOX2/DAPI. **B)** 7 dpi SOX2/MAP2/J2/DAPI. **C)** 14 dpi SOX2/DAPI. **D)** 14 dpi SOX2/MAP2/J2/ DAPI. **E)** 21 dpi SOX2/DAPI. **F)** 21 dpi SOX2/MAP2/J2/DAPI. **G)** Percentages of ventricle cells positive for SOX2 and J2 staining at each timepoint, normalized to the number of SOX2+ cells. **H)** Average nuclear area factor (NAF) at each timepoint.

Organoid measurements and imaging data revealed, especially in the example of African lineage ZIKV infections, smaller organoids and a loss of SOX2 staining (Figs. 2C, 3), suggesting cell loss. ZIKV infection has been shown to cause NPC apoptosis (*43–47*). Because nuclear morphology has been extensively studied as an indicator of apoptosis (*48, 49*), we assessed apoptosis by quantifying nuclear area factor (NAF) values. NAF is calculated as the nuclear area divided by nucleus circularity (*50, 51*), wherein high NAF levels correlate with healthy cells, and very circular nuclei correlate with low NAF values and apoptosis/cell death. At 7 dpi, the comparative differences in NAF values of the seven ZIKV strain infections did not reach statistical significance, relative to Mock, although ZIKV-UG and ZIKV-SEN -infected organoids did have smaller average nuclear areas (Fig. 4H). However, at both 14 and 21 dpi, NAF values for the Asian and American strain ZIKVs (ZIKV-PR, ZIKV-BR, ZIKV-CAM) were similar to Mock, while the SOX2+ cells of African lineage ZIKV-NIV, ZIKV-SEN, and ZIKV-UG infections had significantly lower comparative NAF values. Collectively, the higher NAF values observed with ZIKV-PR, ZIKV-PR and ZIKV-CAM infections suggest differentially low apoptosis as compared to the higher apoptosis in ZIKV-MAL and African lineage infections. The NAF data suggest that the differential organoid pathogenesis that was observed across the three ZIKV lineages (Fig. 4A-F) correlates with degrees of apoptotic cell death.

### African lineage ZIKV and ZIKV-MAL infections disrupt neuronal organization and diminish MAP2 immunofluorescence signal

ZIKV-infected neuronal cell bodies (somata) and dendrites were visualized using an anti-MAP2 antibody (Fig. 5). Although SOX2+ NPCs showed little pathogenesis at 7 dpi in ZIKV-PR, ZIKV-BR, and ZIKV-CAM infections (Fig. 4, rows A and B, columns 2-4), the MAP2+ neuropil was disrupted at 7 dpi in all ZIKV infections (Fig. 5A). In addition, the MAP2 signal (pixels per nucleus) was reduced across all infections (Fig. 5E). At 21 dpi, dsRNA (J2) immunoreactivity was nearly absent from the SOX2+ ventricles of organoids infected by ZIKV-PR, ZIKV-BR, and ZIKV-CAM (Fig. 4, rows A-E and columns 1-4), but anti-dsRNA (J2) signal persisted in extraventricular cells (Fig. 5, row D, columns 2-4). The disruption of the neuropil observed at 7 dpi of the ZIKV-PR, ZIKV-BR, and ZIKV-CAM infections, however, did not progress through 14 dpi and 21 dpi (Fig. 5, rows B and C; Fig. 5E). The neuropil cytoarchitecture in organoids infected by ZIKV-MAL and African lineage viruses was severely disrupted (Fig. 5, rows A-C and columns 5-8), and the corresponding MAP2 area/nucleus decreased precipitously at the 14 dpi and 21 dpi timepoints (Fig. 5E). The disrupted neuropil at 21 dpi is evident in the ZIKV-MAL and African lineage infections (Fig. 5C). Because virus was not cleared from the ventricles of organoids infected with ZIKV-MAL and African lineage ZIKV, the corresponding viral dsRNA (J2) signals from all infected cells, including progenitor cells, are shown in Fig. 5D (columns 5-8). An explanation for these results, similar to that proposed for the SOX2+ progenitor cells, is that Asian and American ZIKV infections retained host antiviral responses, enabling innate immune antiviral responses to modulate virus production. Conversely, a limited host antiviral response in African lineage virus infections could permit persistent virus replication and spread (Fig. 5 row D, columns 5-8).

**Figure 5.**
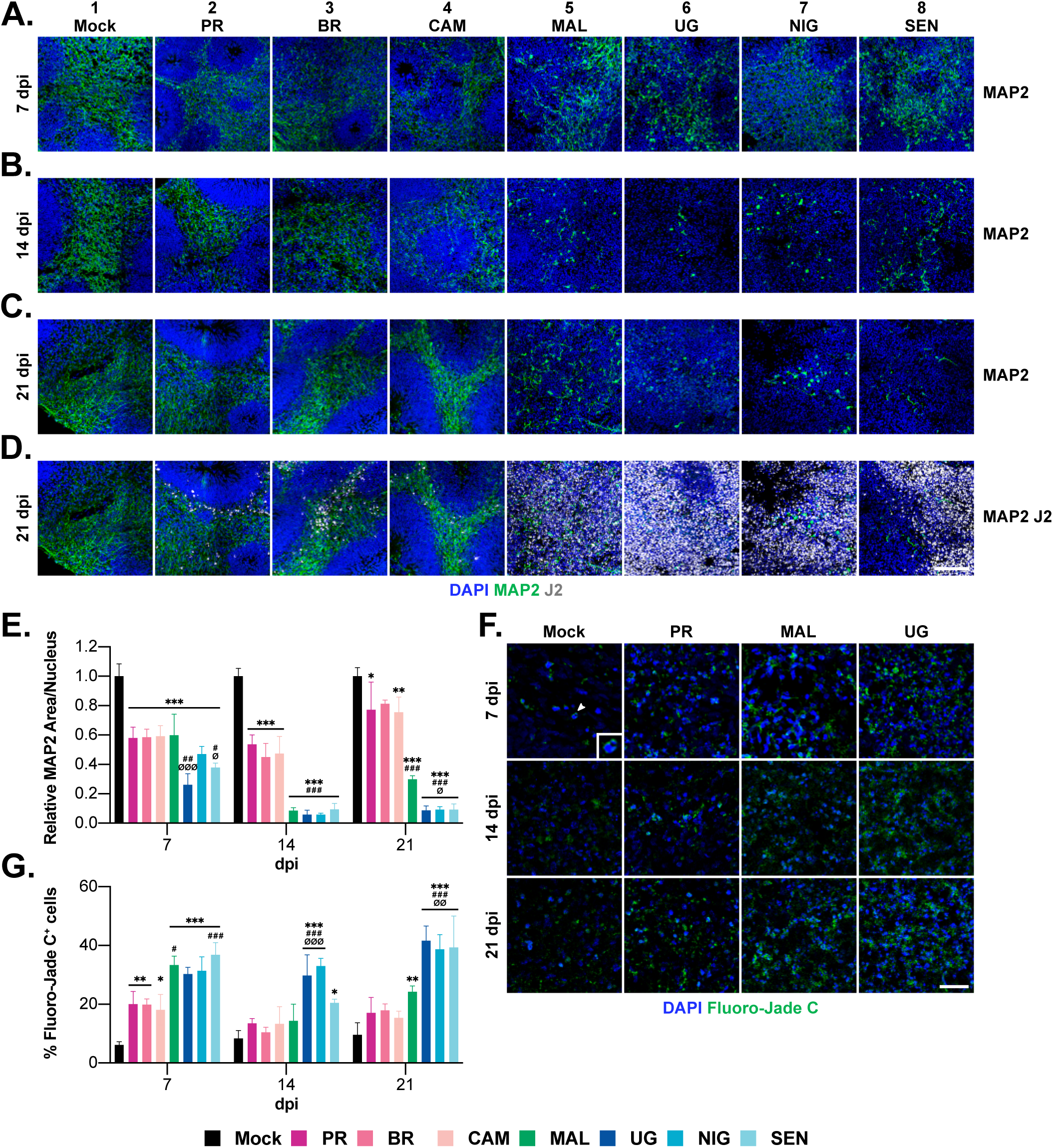
Immunofluorescence microscopy visualizes ZIKV infection in neurons. Confocal microscopy images showing MAP2+ neurons and other cells in the extraventricular regions of human cerebral organoids, stained with anti-MAP2 antibody (green, neuron somata and dendrites), anti-double stranded viral replicative RNA antibody (white, J2), and DAPI (blue) at 7, 14, and 21 dpi (scale bar = 100 μm). The ZIKV strain abbreviations are: PR: Puerto Rico; BRA: Brazil; CAM: Cambodia; MAL: Malaysia; UG: Uganda; NIG: Nigeria; SEN: Senegal. **A-C)** 7, 14, 21 dpi MAP2/DAPI. **D)** 21 dpi MAP2/J2/DAPI. Note that the J2 signal in columns 5-8 includes neurons and NPCs. **E)** Average relative MAP2 area at each timepoint (n = 3, *p ≤ 0.05, **p < 0.01, ***p < 0.001, * compared to Mock, ^#^ compared to PR, BRA, and CAM, ^Ø^ compared to MAL; mean ± SD). **F)** Confocal microscopy images of extraventricular regions stained with Fluoro-Jade C (green) at 7, 14, and 21 dpi in Mock, ZIKV-PR, ZIKV-MAL, and ZIKV-UG infections. The arrow in Mock 7 dpi depicts an FJC+ cell with a zoomed-in version in the lower right corner (scale bar = 25 μm). **G)** Percentages of FJC+ cells at each timepoint (n = 3, *p ≤ 0.05, **p < 0.01, ***p < 0.001, * compared to Mock, ^#^ compared to PR, BRA, and CAM, ^Ø^ compared to MAL; mean ± SD).

Fluoro-Jade C (FJC), a modified Nissl stain, was used to specifically detect degenerating neurons (*52*) (Fig. 5F; quantified in 5G). Compared to Mock, infection by all seven ZIKV strains increased the percentages of FJC+ cells at 7 dpi (Fig. 5G), consistent with the changes in MAP2 area/nucleus (Fig. 5E). Approximately 20% of MAP2+ were FJC+ at 7 dpi in the Asian/American ZIKV infections, while the comparable value for ZIKV-MAL and the African lineage viruses was 30-35%. At 14 dpi and 21 dpi, the number of FJC+ cells in the Asian/American ZIKV infections was not statistically different from mock infections. In most experiments throughout this work, the characteristics of ZIKV-MAL infections mirrored those of the African ZIKVs; however, in the FJC analysis, ZIKV-MAL displayed an intermediate phenotype, showing no increase at 14 dpi, with a small increase at 21 dpi (Fig. 5G). The data show that Asian/American lineage infections caused the lowest neuronal damage, and the African lineage virus infections caused the highest levels of neuronal damage. The FJC staining data strongly suggest that ZIKV infections damage neurons as well as SOX2+ neuronal progenitors, and that the impact is greater in organoids infected by African lineage ZIKV.

### Bulk RNA sequencing reveals differential perturbation of cellular stress gene expression pathways

ZIKV-PR, ZIKV-MAL, and ZIKV-UG, representing American, ancestral Asian, and African lineages, respectively, (*53*) (Fig. 1A), were selected for further study using bulk RNA sequencing (bulk RNA-seq). Bulk RNA-seq reads were mapped to the human reference genome, as well as to the three viral genomes, followed by comparing the transcriptional profiles of mock-infected to virus-infected organoids using cutoff criteria of *p* ≤ 0.05 and absolute fold-change ≥2. Volcano plots derived from virus:mock infection comparisons (Fig. 6 A-C) revealed that the numbers of differentially up-and down-regulated genes at these cutoffs were lowest in the ZIKV-PR infections (Fig. 6A), intermediate in the ZIKV-MAL infections (Fig. 6B), and highest in the ZIKV-UG infections (Fig. 6C), correlating with pathogenesis severity. The 59 upregulated genes in the ZIKV-PR infections (Fig. 6A) include interferon response genes that are in common with the ZIKV-MAL and ZIKV-UG gene expression data. The ZIKV-MAL infection upregulated 450 genes, while ZIKV-UG infection upregulated 564 genes. Following the ZIKV-PR infections, we identified only three downregulated genes meeting cutoff values, whereas 451 and 546 genes were downregulated, respectively, in the ZIKV-MAL and ZIKV-UG infections. These results suggest that pathogenesis severity correlates directly with the numbers of up- and down-regulated genes (*54*) that represent a combination of disease-causing and disease-induced transcripts (*55*).

**Figure 6.**
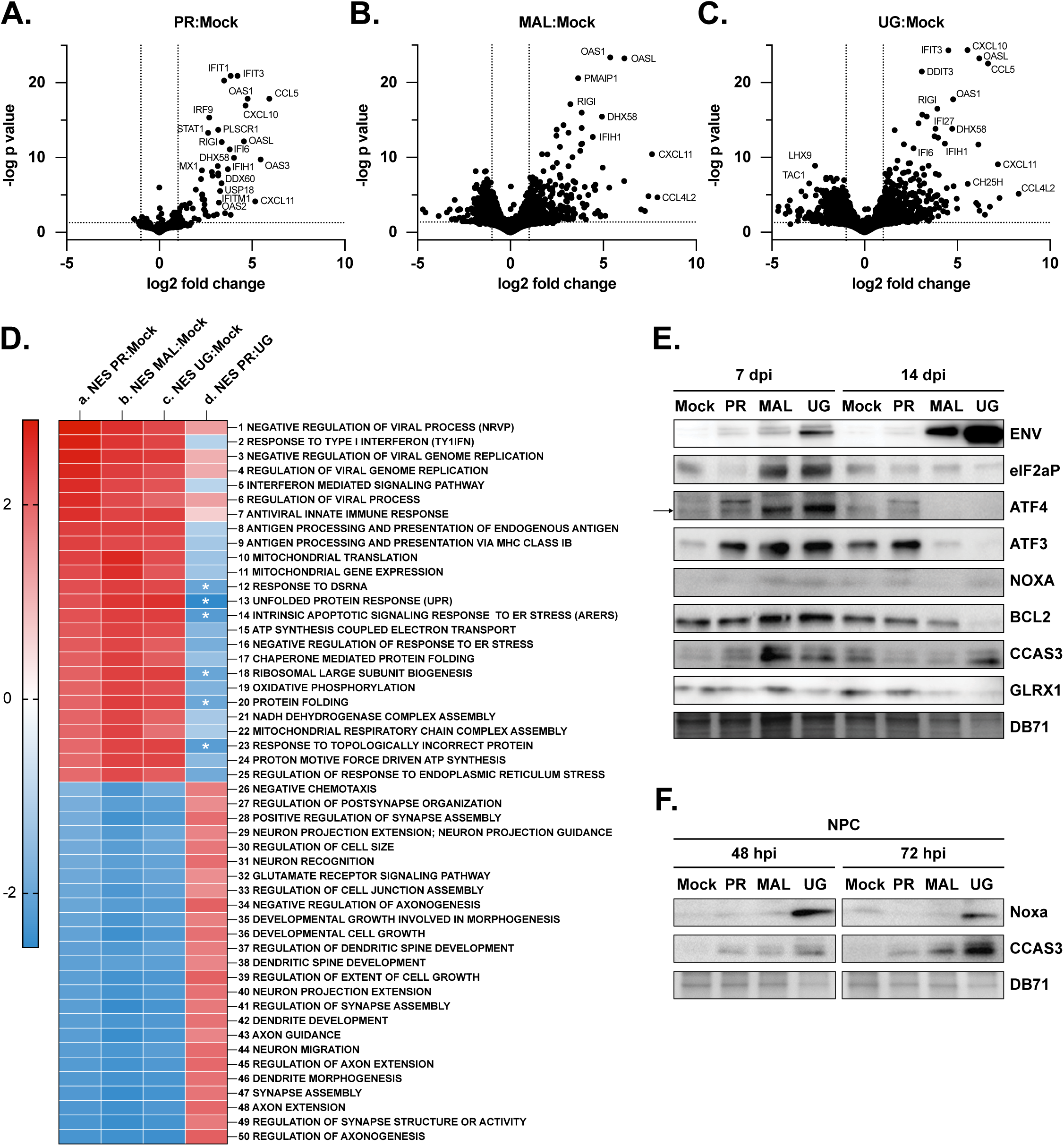
Bulk RNA-seq analysis of cerebral organoids infected with ZIKV-PR, ZIKV-MAL, and ZIKV-UG. **A-C)** Volcano plots showing differentially expressed genes using cutoffs of adjusted p ≤ 0.05 and absolute fold-change ≥2. **D)** Heatmap showing the top 25 positively enriched GSEA pathways (1–25) and the top 25 negatively enriched GSEA pathways (26–50). Column a) ZIKV-PR:Mock NES comparison; Column b) ZIKV-MAL;Mock NES comparison; Column c) ZIKV-UG:Mock NES comparison; Column d) ZIKV-PR:ZIKV-UG NES value comparison. The titles of the GSEA pathways are shown at the right of panel D. All NES values in columns a-c are statistically significant at p ≤ 0.05. Cells indicated by an asterisk (*) in column d are statistically significant at p ≤ 0.05. **E)** Immunoblot showing protein levels of unfolded protein response (UPR) and integrated stress response (ISR) proteins using 7 dpi and 14 dpi organoids infected by ZIKV-PR, ZIKV-MAL, or ZIKV-UG. **F)** Immunoblot showing NOXA (PMAIP) and cleaved caspase 3 (CCAS3) protein in neuronal progenitor cells (NPC) infected by ZIKV-PR, ZIKV-MAL, and ZIKV-UG. The immunoblots are representative of two independent experiments with one technical replicate that sampled three organoids at each time point.

Gene set enrichment analysis (GSEA) was applied to the bulk RNA-seq data as an unbiased approach to evaluate the distribution of gene sets comprising the entire transcriptome, using ranked-ordered differential expression values and statistics, without fold change and *p*-value thresholds. Figure 6D shows a heatmap of normalized enrichment scores (NES) of the top 25 positively-perturbed (columns a-c, rows 1-25) and top 25 negatively-perturbed pathways for ZIKV-PR:Mock, ZIKV-MAL:Mock, and ZIKV-UG:Mock. All of the NES for the positively- and negatively-perturbed pathways in the virus:mock comparisons (columns a-c) are statistically significant (*p* ≤ 0.05). To evaluate differences between ZIKV-PR and ZIKV-UG infections, we analyzed the bulk RNA-seq data as a PR:UG comparison (Fig. 6D, column d). Because of the striking differences in pathogenesis between ZIKV-PR and ZIKV-UG infections (Figs. 3-4), we expected to see corresponding differences in a ZIKV-PR to ZIKV-UG data comparison of antiviral and interferon response GSEA pathways. Antiviral GSEA NES values were positively perturbed in ZIKV-PR infections as compared to ZIKV-UG infections (Fig. 6D, column d, rows 1,3, 4, 6, 12), but only the Response to Double Stranded RNA (DSRNA, row 12) was statistically significant at *p* ≤ 0.05. Two interferon-related pathways, (Fig. 6D, column d, rows 2 and 5) were negatively perturbed in ZIKV-PR infection as compared to ZIKV-UG, suggesting that expression of genes comprising those pathways was downregulated in ZIKV-PR infections as compared to ZIKV-UG, but without statistical significance. Because each GSEA pathway is a composite of many individual genes, we examined a set of twenty-one interferon response genes (ISG) log2 fold change values in a ZIKV-UG to ZIKV-PR comparison. The ISGs showed both up-regulation and down-regulation in the ZIKV-UG to ZIKV-PR comparison (Table 1A); however, there were no statistically significant differences between the two infections, with the exception of downregulated STAT1 in ZIKV-UG infections as compared to ZIKV-PR. Stated another way, STAT1 was upregulated in ZIKV-PR infections over ZIKV-UG. The results of analyzing log2 fold change values for specific ISGs are consistent with the GSEA pathway analysis in confirming that, at the bulk RNA-seq level, ISGs (log2 fold changes) and the NES values for Response to Type 1 interferon and Interferon Mediated Signaling GSEA pathways did not differentiate the ZIKV-PR and ZIKV-UG infections.

**Table 1.** **A)** Log2 fold change and adjusted p-values of interferon stimulated genes (ISG) from a ZIKV-UG:ZIKV-PR comparison. **B)** Log2 fold changes of unfolded protein response (UPR) and integrated stress response (ISR) genes in a ZIKV-UG:ZIKV-PR comparison. The shaded values are statistically significant at p ≤ 0.05. To view a higher resolution image, click here.

| ISG | UG:PR<br>Log2 Fold Change | p adj |
| --- | --- | --- |
| IFIT2 | 0.529 | 0.19158142 |
| IRF7 | 0.22 | 0.247274467 |
| USP18 | 0.115 | 0.656383597 |
| IFIT3 | 0.09 | 0.76810777 |
| IFITM2 | 0.077 | 0.777933385 |
| GBP1 | 0.075 | 0.58864012 |
| MX2 | 0.072 | 0.626820327 |
| IFITM3 | 0.067 | 0.823842043 |
| RSDA2 | 0.025 | 0.884491335 |
| OAS3 | 0.008 | 0.983267382 |
| MX1 | 0.006 | 0.98498309 |
| OAS1 | 0.006 | 0.987206266 |
| OAS2 | -0.015 | 0.944506053 |
| TRIM5 | -0.041 | 0.895239747 |
| IFITM1 | -0.063 | 0.748976106 |
| ISG15 | -0.137 | 0.641143257 |
| IFIT1 | -0.164 | 0.559343926 |
| PKR | -0.37 | 0.278053378 |
| RnaseL | -0.382 | 0.251863411 |
| IRF9 | -0.653 | 0.090170801 |
| STAT2 | -0.672 | 0.098866799 |
| STAT1 | -0.982 | 0.022777503 |

**Table 1. A)** Log2 fold change and adjusted p-values of interferon stimulated genes (ISG) from a ZIKV-UG:ZIKV-PR comparison.
| ISR & UPR<br>Genes | UG:PR<br>Log2 Fold Change | p adj |
| --- | --- | --- |
| DDIT3 | 1.530362102 | 9.75748E-05 |
| PMAIP1 | 1.629868379 | 0.000169666 |
| XBP1 | 1.101738883 | 0.002765331 |
| H2AX | 1.334097259 | 0.004631659 |
| HYOU1 | 1.161220762 | 0.007027641 |
| ATF5 | 1.223028099 | 0.010693759 |
| CHAC1 | 1.409975653 | 0.013870383 |
| PYCR1 | 1.00917709 | 0.026150713 |
| ERp72 | 0.95831037 | 0.034842516 |
| SLC7A11 | 1.46452686 | 0.037900173 |
| TRIB3 | 1.123097412 | 0.040035355 |
| EIF4EBP1 | 0.753796727 | 0.042135097 |
| SHMT2 | 0.710731093 | 0.045830966 |
| PSAT1 | 0.856328086 | 0.072435864 |
| DNAJC3 | 0.917186493 | 0.075308755 |
| ATF3 | 0.965801213 | 0.083920068 |
| PPP1R15A | 0.722884022 | 0.120480025 |
| PDIA6 | 0.48689438 | 0.121403714 |
| HERPUD1 | 0.622020548 | 0.125119146 |
| DNAJB11 | 0.902808308 | 0.17628383 |
| HSP90B1 | 0.324076497 | 0.264328297 |
| HSPA5 | 0.289240927 | 0.291646095 |
| EIF2AK3 | 0.323634018 | 0.329871147 |
| ATF6 | 0.253858549 | 0.346151021 |
| EDEM1 | 0.255185335 | 0.37755941 |
| DNAJB9 | 0.243895856 | 0.435065942 |
| ERN1 | 0.056130871 | 0.820231359 |
| HSPB8 | 0.052797605 | 0.831232288 |

While further comparing ZIKV-UG to ZIKV-PR bulk RNA-seq GSEA data, we found that apoptosis, unfolded protein response (UPR), and integrated stress response (ISR) pathways were positively perturbed with statistical significance in the ZIKV-UG infections as compared to ZIKV-PR infections (Fig. 6D, column d; rows 13,14,16,17, 20, 23, 25). The UPR is activated by the accumulation of unfolded or mis-folded proteins in the endoplasmic reticulum (ER) during high levels of viral protein synthesis in viral replication factories (*56*). The UPR is accompanied by phosphorylation of the alpha subunit of eukaryotic initiation factor (eIF2α) (*57*), causing a shutdown of host protein synthesis as a protective mechanism to conserve energy. Concurrently, eIF2α phosphorylation activates the ISR, causing privileged translation of a set of mRNA transcripts containing upstream open reading frames, including the master regulatory transcription factor ATF4 (*58, 59*). ATF4 upregulates target ISR genes including CHOP (DDIT3) and ATF3 in response to the UPR (*60*). Therefore, to further assess the GSEA NES values (Fig. 6), log2 fold change values for specific ISR and UPR genes are presented in Table 1B. These data agree with the pathway analysis in demonstrating that many ISR and UPR genes were positively perturbed with significance (green shading) in the ZIKV-UG infections as compared to ZIKV-PR infections. The results strongly suggest that stress responses are elevated in ZIKV-UG infections as compared to ZIKV-PR infections, directly correlating stress responses and the severe pathogenesis observed in ZIKV-UG infections (Figs. 3-4).

To validate the ISR/UPR transcriptome signature, immunoblotting was performed to define the translated cellular stress response protein levels. The ZIKV viral envelope protein (ENV) was detected as a measure of virus levels at 7 dpi and 14 dpi (Fig. 6E). eIF2α phosphorylation is a marker of cell stress, and both ZIKV-MAL and ZIKV-UG -infected organoids showed increased levels of phosphorylated eIF2α relative to mock-infected cells at 7 dpi. On the contrary, ZIKV-PR infection did not increase the levels of phosphorylated eiF2α relative to mock-infected organoids (Fig. 6E). The eiF2α phosphorylation signal was greatly diminished by the 14 dpi time point, suggesting a transient response, or that the signal diminished because of cell death. ATF4 levels increased in the ZIKV-MAL and ZIKV-UG infections. ATF4 RNA is translationally controlled (*58*) and ATF4 protein controls expression of ATF3, also a transcription factor that can be functionally pro-apoptotic or adaptive (*61*). Immunoblotting revealed that ATF3 and ATF4 proteins were expressed at higher levels in the ZIKV-MAL and ZIKV-UG infections at 7 dpi, later diminishing by 14 dpi in the same infections (Fig. 6E). Elevated ATF3 and ATF4 levels are markers of integrated stress responses. NOXA (PMAIP) is an ATF4-induced pro-apoptotic protein that is elevated in the 7 dpi ZIKV-UG organoid infections (Fig. 6E) and also confirmed in NPC infections (Fig. 6F), suggesting increased apoptotic activity in the ZIKV-UG infections. The anti-apoptotic BCL2 protein signal is comparable among all infections at 7 dpi (Fig. 6E), but the levels in 14 dpi ZIKV-UG infections were low compared to ZIKV-PR, suggesting diminished anti-apoptotic signaling in response to ZIKV-UG infections. Apoptosis was suggested by cleaved caspase 3 (CCAS3) levels, which were higher in ZIKV-MAL and ZIKV-UG than ZIKV-PR infections in organoid immunoblots (Fig. 6E), as well as NPC immunoblots (Fig. 6F). These data suggest relatively low apoptotic activity in the ZIKV-PR infections, higher levels in the ZIKV-MAL infections, and highest levels in ZIKV-UG infections at 14 dpi. Finally, we examined glutaredoxin 1 (GLRX1) protein, which contributes positively to the antioxidant defense system (*62*). GLRX1 levels are reduced in ZIKV-UG infections at both 7 and 14 dpi, and in ZIKV-MAL infections at 14 dpi, suggesting greater protection against oxidative stress in the ZIKV-PR infections (*63*). The results presented in Fig. 6E-F are consistent with the bulk RNA-seq data, suggesting that elevated stress responses in the ZIKV-MAL and ZIKV-UG infections were accompanied by UPR and ISR activation (*64*). The elevated stress responses in ZIKV-MAL and ZIKV-UG correlate with the corresponding severe pathogenesis observed in the corresponding immunofluorescence imaging (Figs. 3-4).

### Single cell RNA sequencing identifies cell types and differential gene expression pathways

Single cell RNA sequencing (scRNA-seq) was used to identify organoid cell types, their infection status, and their transcriptome changes following virus infections. Nine cell clusters were identified by marker expression (Fig. 7 A-B). Viral RNA expression quantification in the clusters (Fig. 7C-E) demonstrates that all cell types were infectable by the three ZIKV lineages, and that the viral RNA expression levels were similar, varying by a factor of approximately two-to-four at 7 dpi, without evidence of greater tropism for one cell type over another. The named clusters are: neuronal progenitor cells (NPC), intermediate progenitor cells (IPC), neurons, midbrain-hindbrain (MB:HB) cells, ISG-High cells (*65*), NPC-like cells, and astrocytes. The NPC-like cells and the ISG-High cells did not clearly match known cell type gene expression profiles, suggesting that they may be in intermediate development differentiation stages or represent multiple cell types whose clustering was driven by ISG transcript levels, respectively. The numbers of cells analyzed from each cluster are shown in Table S1.

**Figure 7.**
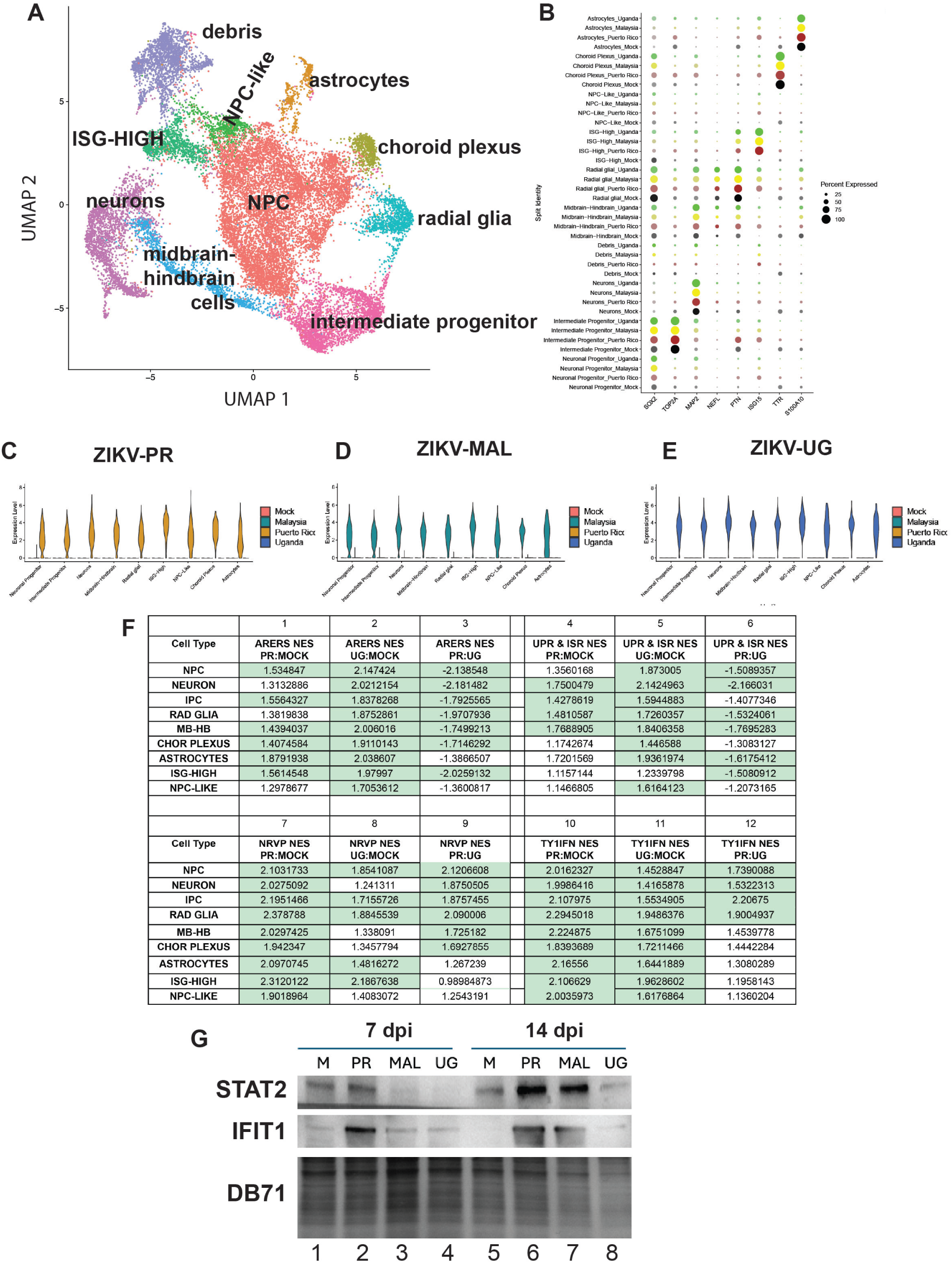
Single cell RNA-seq reveals differential pathway enrichments that distinguish cell types and ZIKV lineages. **A)** UMAP clustering of single cell RNA-seq data. NPC: neuronal progenitor cell; ISG: interferon stimulated gene. **B)** Bubble plot showing marker expression data used to assign cell types. **C-E)** Viral RNA expression in infected organoids at 7 dpi, grouped by ZIKV lineage. **F)** NES values of ZIKV-PR:Mock, ZIKV-UG:Mock, and ZIKV-PR:ZIKV-UG for the response to ER stress (ARERS), unfolded protein response/integrated stress response (UPR/ISR), response to negative regulation of viral process (NRVP), and response to type 1 interferon (T1IFN) GSEA pathways. Shaded NES values are statistically significant at p ≤ 0.05. **G)** Immunoblot analysis showing STAT2 and IFIT1 protein levels in organoids infected by ZIKV-PR, ZIKV-MAL and ZIKV-UG. DB71 dye was to assess protein loading.

#### Stress and apoptosis responses

To assess stress responses in the ZIKV-infected organoids, virus:mock and virus:virus comparisons of scRNA-seq NES values were run for Apoptotic Response to Endoplasmic Reticulum Stress (ARERS) and Unfolded Protein Response (UPR) pathways in all cell cluster types (Fig. 7F). Virus:mock comparisons are shown in columns 1-2 and 4-5, respectively, while the corresponding PR:UG virus:virus comparisons are presented in columns 3 and 6). The PR:Mock infection data (column 1) suggest that ZIKV-PR infection led to significant positive perturbation of apoptosis genes in the ARERS GSEA pathway in most cell types; however, the NES values were not significantly higher or lower as compared to Mock in neurons, radial glia, or NPC-like cells. Conversely, the ZIKV-UG:Mock infection was accompanied by higher NES values, relative to ZIKV-PR:Mock, which were all statistically significant (column 2). When the NES values were compared as virus:virus (ZIKV-PR to ZIKV-UG), the ARERS NES values are negatively perturbed in ZIKV-PR infections relative to ZIKV-UG (column 3) in all cell types but astrocytes and NPC-like cells. These data suggest that ZIKV-PR infections induce lower stress responses as compared to ZIKV-UG infections.

We next examined the unfolded protein response GSEA pathway, which also includes integrated stress response genes. Similar to the ARERS results, the ZIKV-PR infection positively perturbed UPR pathway genes overall, but the NES values were significantly different from Mock only in neurons, IPC, radial glia, and midbrain-hindbrain cells (Fig. 7F, column 4). The ZIKV-UG infection positively perturbed UPR pathway genes (column 5) to levels higher than ZIKV-PR (column 3), and with statistical significance in all cell types but the ISG-High. In the virus:virus ZIKV-PR to ZIKV-UG comparison, UPR pathway genes were uniformly negatively perturbed, reaching statistical significance in all but IPC, choroid plexus, and NPC-like cells (Fig. 7F, column 6). Taken together with the ARERS results (column 3), the data strongly suggest that ZIKV-PR infections resulted in low stress (ARERS and ISR/UPR) and mild pathogenesis, while the ZIKV-UG infections resulted in high stress and severe pathogenesis (Figs. 3-4).

#### Antiviral and interferon response pathways analysis

ZIKV-UG pathogenesis is severe in comparison to the mild ZIKV-PR pathogenesis (Figs. 3-4); however, the bulk RNA-seq (ZIKV-PR:ZIKV-UG) analysis (Fig. 6D, column d, rows 1-7, 12) showed that the comparative antiviral (NRVP) and type 1 interferon (TY1IFN) pathway NES values were statistically indistinguishable (*p* ≥ 0.05), with the exception of the Response to DSRNA. Of note, double stranded RNA is expressed as a replicative intermediate form during viral RNA replication and is sensed by RIG-I. Given the striking differences in pathogenesis, it was somewhat surprising that the antiviral and type 1 interferon response NES values were not statistically distinguishable in comparing the ZIKV-PR and ZIKV-UG infections. We therefore turned to single cell RNA-seq data to further explore antiviral NRVP NES responses across the cell types in both virus:mock and virus:virus comparisons. NES values from ZIKV-PR:Mock comparative scRNA-seq data were uniformly positively perturbed and statistically significant (Fig. 7F, column 7). The comparable NES values from the ZIKV-UG:Mock comparison were lower than ZIKV-PR values (Fig. 7F; compare columns 7 and 8), and the NES values failed to reach significance (*p* ≥ 0.05) in comparison to mock infected neurons, midbrain-hindbrain, choroid plexus, and NPC-like cells (column 8). The ZIKV-PR:ZIKV-UG comparison revealed that the NRVP antiviral pathway was significantly positively perturbed in ZIKV-PR infections, as compared to ZIKV-UG, in all cell types but astrocytes, ISG-High, and NPC-like cells (column 9). These data strongly suggest that ZIKV-PR infections induced higher levels of antiviral gene pathway expression responses (NES values) in some cell types, as compared to ZIKV-UG infections. The higher level antiviral NRVP expression in the ZIKV-PR infection therefore correlates with mild pathogenesis, while the converse was true for the ZIKV-UG infections (Figs. 3-4).

In parallel, interferon responses were compared for the ZIKV-PR:Mock infection via NES values of the Response to Type 1 interferon pathway (TY1IFN). The ZIKV-PR:Mock comparison showed positive perturbation with significance in all cell types (Fig. 7F, column 10), with higher NES values than observed in the ZIKV-UG:Mock infection (Fig. 7F, column 11). The results of the ZIKV-PR:ZIKV-UG comparisons demonstrate that the NES values are positively perturbed in the ZIKV-PR infection, relative to ZIKV-UG, with significance in NPC, neurons, IPC, and radial glia (Fig. 7F, column 12). These four cell types are present in organoid ventricles which are surrounded by newly born neurons (Fig. S1). These data suggest that the NRVP and TY1IFN pathway responses are induced to higher levels in the ZIKV-PR infections, as compared to ZIKV-UG, in neurons and in ventricle cells (NPC, IPC, radial glia). This enhanced response to type 1 interferon marks a distinction between ZIKV-PR and ZIKV-UG infections in some cell types, and correlates with the clearing of American lineage ZIKV, but not African lineage ZIKV from ventricles (Fig. 4).Therefore, the positive perturbation of antiviral pathways in ventricle cells and neurons could create an antiviral environment that clears American lineages viruses from ventricles, as shown by the dramatic decrease in SOX2+/J2+ cells at 14 dpi (Fig. 4G).

ISG log2 fold changes were found to be statistically indistinguishable in the ZIKV-PR and ZIKV-UG infections, with the exception of STAT1 (Table 1A), leading us to question how to reconcile the differential pathogenesis (Figs. 3-4) with similar ISG expression levels. ISG expression is activated by interferon binding to interferon receptors, which signal through JAK-STAT proteins to further activate transcription factors that bind the interferon sensitive response element (ISRE) for ISG expression. It was reported previously that STAT2 protein is degraded in Dengue and Zika virus infections, facilitated by non-structural protein 5 (NS5) (*22, 66*). We therefore used immunoblotting to examine STAT2 protein levels. The results revealed detectable STAT2 protein in organoid samples from ZIKV-PR infections (Fig. 7G, lanes 2 and 6); however, STAT2 protein levels were diminished in the ZIKV-MAL and ZIKV-UG infections at both 7 dpi and 14 dpi (Fig. 7G, lanes 3, 4, 7, 8). We further examined IFIT1, which is an interferon-stimulated antiviral protein and a target of JAK-STAT signaling. Similar to STAT2, IFIT1 levels were detectable in ZIKV-PR infections at both 7 and 14 dpi (Fig. 7G, lanes 2 and 6), while diminished in the ZIKV-MAL and ZIKV-UG infections (lanes 3, 4, 7, 8). These results suggest that STAT2 protein was degraded, as observed in Dengue virus infections, which would be predicted to cause a signaling defect in the ISG expression pathway in ZIKV-MAL and ZIKV-UG infections. The presence of STAT2 protein in ZIKV-PR infections would coincide with ISG expression and antiviral responses, while attenuated antiviral responses in ZIKV-MAL and ZIKV-UG infections would limit the antiviral response. The STAT2 and IFIT1 data (Fig. 7G) support a hypothesis stating, first, that host antiviral responses to the ZIKV-PR infection were intact, creating an antiviral environment that cleared virus from organoid ventricles (Fig. 4G). Second, antiviral responses were limited in the ZIKV-MAL and ZIKV-UG infections due to diminished STAT2 levels, thereby permitting ongoing virus replication and spread (Fig. 4, rows D and F).

### ZIKV-UG infection disrupts the balance between antioxidants and reactive oxygen species, contributing to pathogenesis

Several lines of evidence reported here, including bulk and single cell RNA sequencing, immunoblotting, and imaging pointed to stress as a pathogenesis mechanism in the ZIKV infections. Stress responses in viral infection can correlate with ROS imbalance (*67–70*); therefore, to evaluate possible ROS imbalance in ZIKV infections, we used ZIKV-infected Vero cells and MitoSOX dye to detect mitochondrial superoxide production. At 24 hours post-ZIKV-PR infection, few cells exhibited a positive ROS signal (Fig. 8A). In contrast, ROS levels were elevated in cells infected by ZIKV-MAL and ZIKV-UG, with ZIKV-UG showing the highest MitoSOX signal (Fig. 8A-B). ZIKV-UG infection not only correlated with the highest number of MitoSOX-positive cells (Fig. 8B) but also with greater average fluorescence intensity per cell compared to ZIKV-PR or ZIKV-MAL infections (Fig. 8C). These results demonstrate that mitochondrial superoxide levels increased in ZIKV-MAL and ZIKV-UG-infected Vero cells.

**Figure 8.**
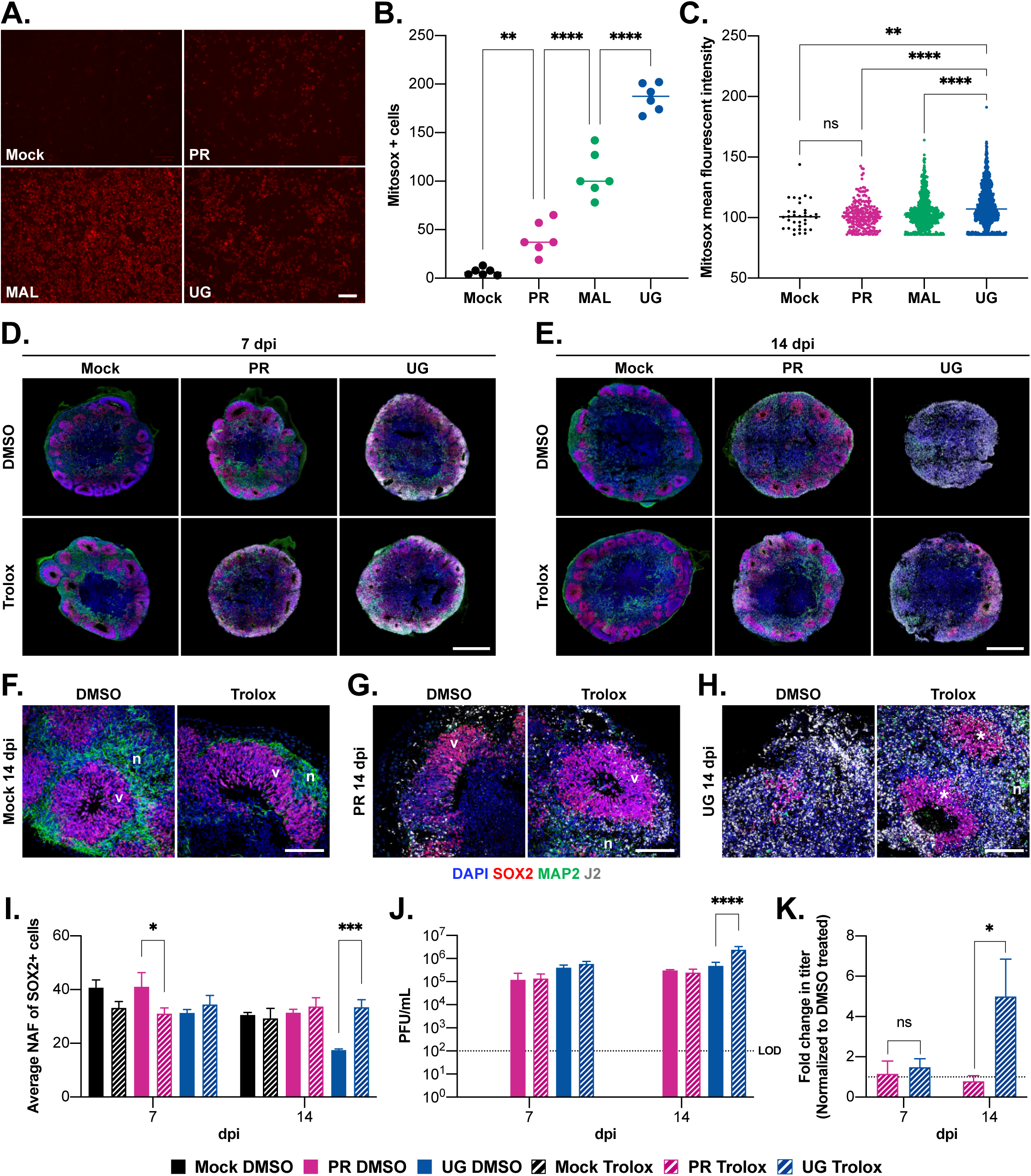
Trolox, a hydroxyl radical scavenger, partially protects ZIKV-UG-infected cerebral organoids from severe pathogenesis. **A)** Visualization of superoxide production in ZIKV-infected Vero cells using MitoSOX Red dye (scale bar = 100 μm). **B-C)** quantification of ROS+ cells and ROS fluorescence intensity per cell for each of the ZIKV infections shown in Panel A. **D)** Confocal immunofluorescence microscopy of cerebral organoids at 7 dpi with ZIKV-PR or ZIKV-UG and treated with the dimethyl sulfoxide (DMSO) carrier or DMSO containing 100 μM Trolox (scale bar = 500 μm). **E)** Same as D at 14 dpi. **F-H)** Zoomed-in views of organoids that were mock-infected (F), ZIKV-PR-infected (G) or ZIKV-UG infected (H) and treated with either DMSO or Trolox. n: MAP2-stained neuron; v: SOX2-stained ventricle. The arrow in panel H shows remnant SOX2+ cells following ZIKV-UG infection. Asterisks (*) in panel H mark ventricles that are spared by Trolox treatment from ZIKV-UG infection-mediated cytoarchitecture disruption (scale bar = 100 μm). **I)** NAF analysis of images in panels D-E to assess apoptosis at 7 dpi and 14 dpi, with and without Trolox treatment. **J)** Viral titers representing 24 hrs of released virus at 7 and 14 dpi in Trolox treated and control organoids **K)** Fold change in viral titers at 7 and 14 dpi following resulting from Trolox treatment.

To directly test ROS imbalance as a mechanism underlying differential ZIKV pathogenesis, we used Trolox, a vitamin E analog that scavenges superoxides and prevents oxidative stress-induced apoptosis (*71, 72*). Organoids were treated with either Trolox or DMSO carrier following ZIKV-PR or ZIKV-UG infection, and processed for immunofluorescence microscopy to assess cytoarchitecture, as well as apoptosis using NAF values. At both 7 dpi (Fig. 8D) and 14 dpi (Fig. 8E), the mock-infected DMSO-treated control organoids maintained SOX2+ ventricles with clear MAP2 neuronal soma and dendrite staining, indicating intact cytoarchitecture.

Similarly, mock-infected organoids treated with Trolox retained their ventricle cytoarchitecture. By 14 dpi, organoids infected with ZIKV-PR retained organized ventricles, and Trolox treatment did not induce any notable morphological changes (Fig. 8E). However, Trolox had a profound effect on 14 dpi ZIKV-UG-infected organoids, wherein, instead of the severe pathogenesis (Fig. 4), we observed organized ventricles and SOX2+ cells (Fig. 8E). Higher magnification images of the Trolox treatments are shown in Fig. 8F-H. At 14 dpi, the mock-infected and Trolox-treated organoids retained organized SOX2+ ventricles (v) and MAP2+ neurons (n) (Fig. 8F), similar to the appearance of ZIKV-PR-infected organoids (Fig. 8G). We noted earlier (Fig. 4) that in the 14 dpi ZIKV-PR infections, the SOX2+ cells present in ventricles have diminished dsRNA (J2) signal as compared to surrounding areas or the ZIKV-UG infection, suggesting that ZIKV-PR infection in NPC resolves by 14 dpi (Fig. 4, Fig. 8G). At 14 dpi, ZIKV-UG infection disrupted ventricle cytoarchitecture and diminished SOX2 staining in the DMSO control (Fig. 8H, left image); however, in the presence of Trolox (Fig. 8H, right image), organized ventricles are evident, including SOX2+ cells with decreased J2 reactivity (labeled *) and MAP2+ neurons (n).

We further hypothesized that the improved ventricle cytoarchitecture observed with Trolox treatment might correlate with reduced apoptosis. Apoptosis was assessed by NAF analysis. Trolox treatment of ZIKV-PR-infected organoids at 7 dpi led to a slight pro-apoptotic effect (lower NAF values) (Fig. 8I). This effect was unexpected; it may be Trolox-related because Trolox has been reported to induce pro-apoptotic activity under certain conditions (*73*). A large increase in NAF values, indicating reduced apoptosis, was observed in Trolox-treated ZIKV-UG-infected organoids at 14 dpi (Fig. 8I). These results suggest that ROS-induced apoptosis contributes to the observed disruption of cytoarchitecture and loss of SOX2+ cells during ZIKV-UG infection.

As a further consideration of regulatory mechanisms, we considered that Trolox’s ROS-scavenging activity reduced pathogenesis in the ZIKV-UG-infected organoids by either a) reducing cell death by decreasing apoptosis or b) lowering virus replication and virus spread, thereby maintaining SOX2+ progenitor cells and cytoarchitecture. Significant changes in viral titers were not observed at 7 dpi (Fig. 8J); however, at 14 dpi, the ZIKV-UG infection titer increased five-fold with Trolox treatment (Fig. 8J-K). These data suggest that the improved cytoarchitecture results from diminished ROS-induced apoptosis, and that ZIKV-UG viral titer is not wholly causal for ZIKV pathogenesis in cerebral organoids derived from human embryonic stem cells.

## DISCUSSION

Zika virus (ZIKV), recognized for its pandemic potential by the World Health Organization, is a significant human pathogen. Differential pathogenesis among Zika virus lineages and strains has been described (*74–77*); however, the underlying mechanisms have not been elucidated. Because viral pathogenesis is a complex, multifactorial process, we have studied ZIKV pathogenesis by comparing infections caused by African, Asian, and American lineages, reasoning that differences and similarities among the infections and their pathogenesis properties may illuminate causative mechanisms.

Imaging data revealed differential pathogenesis when comparing African and Asian/American ZIKV infections (Figs. 3-4). In the Asian/American ZIKV infections, but not in ZIKV-MAL or African lineage virus infections, double stranded replicative intermediate RNA levels were greatly reduced in the ventricles by 14 dpi (Fig. 4G). One interpretation of these results is that by 14 dpi, ventricle cells in Asian/American ZIKV infections had differentially established an antiviral environment that reduced virus replication and spread, resulting in virus clearance. Conversely, the ventricle cells infected by ZIKV-MAL and African lineage ZIKV-UG did not establish an antiviral environment, thereby permitting unchecked virus replication and apoptotic NPC death (*43*). We expected that the antiviral responses would include robust differential ISG induction. The STAT2 protein loss (Fig. 7G) in ZIKV-MAL and ZIKV-UG infections would be expected to prevent signaling to induce ISG transcription from the interferon-stimulated regulatory element (ISRE). However, bulk RNA-seq data did not identify significant differences in either the log2 fold ISG expression values or the GSEA NES values of interferon and antiviral response pathways when comparing ZIKV-PR to ZIKV-UG infections (Fig. 6A-C; Table 1A; Fig. 6D, columns d and e, rows 1-7). The absence of differential ISG transcription in the ZIKV-PR and ZIKV-UG infections, considering the contrasting pathogenesis severities (Figs. 3-4) suggested either that mechanisms other than STAT2-mediated ISG expression defined differential pathogenesis, or alternatively that the bulk RNA-seq approach did not capture key mechanistic details.

We considered that, by processing organoids *en masse* for bulk RNA-seq, the data averaging over heterogeneous cells may have masked evidence of cell-specific gene expression (*78, 79*). scRNA-seq approaches compared the GSEA NES values for individual organoid cell types (Fig. 7A) in the context of antiviral (NRVP) and interferon (TY1IFN) response pathways (Fig. 7F; columns 7-9 and 10-12). In contrast to the bulk RNA-seq data, scRNA-seq revealed that interferon and antiviral pathway responses could be distinguished in the ZIKV-PR and ZIKV-UG infections, showing positive perturbation with statistical significance in ZIKV-PR infections of neurons, NPC, radial glia, and IPC. This result suggests that important gene expression data were masked in bulk RNA-seq, and that differential antiviral response in ventricle cells and neurons could explain the limited viral replication and spread observed in ZIKV-PR infections (Figs. 3-4). Conversely, the negatively perturbed antiviral and interferon response pathways, as well as STAT2 protein degradation would create an environment favorable to unchecked viral RNA replication and spread in ZIKV-UG infections.

Returning to the bulk RNA-seq data, we identified statistically significant differential positive perturbation of host stress response pathways in ZIKV-MAL and ZIKV-UG infections, as compared to ZIKV-PR infections. These pathways include apoptosis, oxidative phosphorylation, UPR, and ISR (Fig. 6D, columns d-e, rows 12-13, 15, 17, 21). Both UPR and ISR stress responses center on eIF2α phosphorylation, which slows or blocks cellular protein synthesis and activates the integrated stress response (ISR) (*59*). The ISR is activated by signaling to phosphorylate eIF2α, followed by transcription and translation of the master regulator, ATF4, to enable preferential translation of messenger RNAs with open reading frames (uORFs) that are upstream of the canonical AUG initiation codon. The preferentially translated mRNAs include amino acid biosynthesis and transport proteins, antioxidant defense proteins, ER stress response proteins, and apoptosis-causing proteins that appear in prolonged stress (*80–82*) (Fig. 6E). Our results (Fig. 6D-E) demonstrate that organoids infected with the African lineage viruses differentially activated UPR/ISR cellular responses, which can be adaptive for cell survival (ZIKV-PR) infections, or maladaptive for apoptotic cell death (ZIKV-MAL and ZIKV-UG infections) (*59, 83, 84*). The data presented in this study strongly suggest that ISR hyperactivation in severe ZIKV-UG infections led to DDIT3-NOXA mediated apoptosis (*85*), while conversely, moderate ISR activation in ZIKV-PR infections was adaptive and therefore promoted cell survival.

Other experiments showed that ZIKV-MAL and ZIKV-UG infections were characterized by high levels of superoxides (Fig. 8A-C), correlating ROS imbalance to ZIKV pathogenesis (*68, 69, 86, 87*). ROS imbalance has myriad potential effects (*88*), including cytochrome C release that activates the caspase apoptosis cascade or causes direct damage to DNA, proteins, or lipids. We interpret the results of the Trolox experiments (Fig. 8) as evidence that ROS imbalance is a critical mechanism of ZIKV pathogenesis. We note that a recent publication (*89*) concluded that ROS is not a significant pathogenesis determinant for ZIKV infections in SH-SY5Y cells. An important distinction is that we tested both African and Asian lineage ZIKV, while Mendonça-Vieire *et al*. (*89*) limited their analysis to an American lineage ZIKV (Brazil, PE243), and did not test African lineage ZIKV. In fact, our results agree with Mendonça-Vieira *et al*. in the context of Asian/American lineage viruses, where there was limited evidence for elevated ROS-related pathogenesis, but clear evidence for ROS imbalance in the African lineage ZIKV infections.

One potential mechanism of the partial rescue of organoid cytoarchitecture and neuronal progenitor cells by Trolox is that the treatment was antiviral and reduced viral RNA replication. Quantifying infectious virus release revealed that, instead of diminished viral RNA production, viral titer increased following Trolox treatment (Fig. 8J-K). Trolox treatment was accompanied by decreased apoptosis (Fig. 8I) that would improve organoid health and homeostasis, potentially creating a favorable environment for ongoing virus replication. However, higher ZIKV titer, paired with diminished apoptosis and improved organoid cytoarchitecture, is somewhat counterintuitive. Yockey *et al.* (*90*) reported related findings, however, by demonstrating that ZIKV-infected IFNAR-/- mouse pups had significantly better morbidity/mortality outcomes than IFNAR+/+ mice, despite the fact that Zika virus replicated to nearly 100-fold higher levels in the IFNAR-/- mouse pups. Our results parallel others (*90, 91*) in demonstrating that virus titer does not always correlate directly with pathogenesis, emphasizing that pathogenesis is defined by host responses to the virus infection.

The diminished STAT2 protein levels in ZIKV-MAL and ZIKV-UG infections (Fig. 7G) are consistent with Grant *et al.* (*22*), who reported that STAT2 is more stable in ZIKV-PR infections (Vero cells) than ZIKV-UG infections, where STAT2 was degraded by the proteasome. In Dengue virus infections, STAT2 is marked for proteasomal degradation by a ubiquitin ligase (UBR4) that is recruited by the viral non-structural protein 5 (NS5) (*92*). However, ZIKV infections do not require the UBR4 E3 ubiquitin ligase to induce STAT2 degradation (*22*). One hypothesis to explain the differential STAT2 degradation is that the ZIKV-PR NS5 protein differentially facilitates proteolysis among the lineages, perhaps because of amino acid substitutions that distinguish the two lineages. Further research to define mechanisms of STAT2 degradation in ZIKV is needed to understand the correlation with pathogenesis.

Asian/American lineage ZIKV infections can cause Congenital Zika Syndrome (*93*), which includes microcephaly. Although the American/Asian ZIKV studied here showed relatively mild pathogenesis compared to African lineage ZIKV, the ZIKV-PR strain does indeed cause organoid pathology. Our data are consistent with Tripathi *et al.* (*91*), who concluded that African lineage ZIKV cause a short, severe pathogenesis associated with fetal demise in a mouse model, while the Asian/American ZIKV infections cause a prolonged period to pathogenesis, occasionally causing death. Other data suggest that synapse impairment is a feature of CZS (*94*). The comparative MAP2 imaging data (Figs. 3, 5) demonstrate that neuronal structure is disrupted in ZIKV infections; moreover, bulk RNA-seq data demonstrate that many gene pathways related to neuronal development are downregulated in ZIKV-MAL and ZIKV-UG infections, including neuronal migration and axonal guidance genes (Fig. 6D, shaded blue).

Although cerebral organoids derived from human embryonic stem cells have many advantages for studying virus infections (*36*), a limitation is that organoids lack vasculature and are therefore limited by diffusion, which can lead to a necrotic core. We used small and early organoids, maximally approximately two millimeters in diameter, and only 21 days post-embryoid body formation to minimize the necrotic core. A second potential limitation of the study is that immune cells are not present in the cerebral organoid cultures. Organoid technology continues to advance through co-culture with immune and glia cells (*95, 96*)). In this study, the comparative analysis provided new insights into innate immune responses that differentiated the pathogenesis caused by Zika virus strains.

Areas for future research into effects of ZIKV infections on axonal guidance and neuronal migration could include studies of extracellular matrix glycoproteins such as reelin, slit, and semaphorins, as well as their receptors, robo and plexin, which regulate neuronal guidance migration during brain development. Another potential area of further research, based on the partial restoration of organoid cytoarchitecture by the ROS scavenger, Trolox, would be to consider antioxidants as therapeutics for ZIKA disease (*97*). To date, the success of antioxidant therapies has been mixed (*98*), but their potential value has been emphasized (*88, 99*), (*100*). Overall, side-by-side comparisons of virus infections in cerebral organoids will have ongoing value in defining additional pathogenesis mechanisms toward designing potential therapeutic approaches.

## MATERIALS AND METHODS

### Study Design

The objective of this study was to understand how infections caused by closely related viruses can result in diametrically opposed pathogenesis severities. We studied Zika viruses (ZIKV), which have pandemic potential as well as causing serious neurological sequelae, such as Guillain-Barré syndrome and Congenital Zika Syndrome (CZS). The rationale of the study was that, by analyzing side-by-side comparative infections using ZIKVs that cause mild or severe pathogenesis, mechanistic details would be revealed to define the molecular basis of differential pathogenesis. The experiments were performed in a relevant human mode system, using organoids derived from human embryonic stem cells at a time corresponding to the first trimester of human development. The organoids were infected at a very low multiplicity of infection, mimicking comparable amounts of virus delivered by a mosquito bite, and allowing us to follow the infections over a period of up to 21 days. The data revealed that African lineage ZIKV caused severe pathogenesis, while American and Asian lineage ZIKV caused relatively mild pathogenesis. Bulk RNA sequencing was used to examine transcriptomic changes associated with ZIKV infections and demonstrated that stress response pathways (unfolded protein response and integrated stress response) are differentially upregulated in African lineage infections as compared to Asian/American lineage ZIKV, correlating with severe pathogenesis. Single cell RNA sequencing demonstrated that all nine cell types were infected by ZIKV, without revealing a tropism for one particular cell type. Single cell RNA-seq confirmed that stress response pathways were upregulated in most cell types following African lineage ZIKV infections and also demonstrated that antiviral and interferon pathway genes were differentially upregulated in Asian/American lineage ZIKV, offering a potential explanation for why ZIKV replication, spread, and pathogenesis were limited in American lineage ZIKV infections. Polyacrylamide gel electrophoresis and immunoblotting were used to examine levels of stress-response proteins that accompanied mild and severe pathogenesis. In addition, we found that the STAT2 protein, which is critical for signal transduction leading to the expression of antiviral interferon-stimulated genes (ISG), is differentially degraded in African lineage infections, limiting antiviral ISG expression and enabling unfettered African lineage ZIKV replication and spread. Cell staining to detect hydroxyl radicals was performed, revealing that African lineage ZIKV infections are characterized by high reactive oxygen species (ROS) levels. Trolox, a hydroxyl radical scavenger, was found to partially block pathogenesis caused by African lineage ZIKV, suggesting a potential therapeutic approach for treating ZIKV patients. An NIH-registered stem cell line was used for all experiments, following protocols monitored by the Committee on Assessment of Biohazards and Embryonic Stem Cell Research Oversight at the Massachusetts Institute of Technology.

### NGD 0.5X media

NGD 0.5X media was prepared as previously described (*101*). 500 ml of NGD 0.5X consists of a mixture of 475 ml Neurobasal media (Thermo), 5 ml Gem21-VitA (0.5X, Gemini Bioproducts), 2.5 ml Neuroplex N2 (0.5X, Gemini Bioproducts), 5 ml pyruvate (100 mM), 5 ml Glutamax, 5 ml penicillin/streptomycin, 5 ml NaCl (5M), 1 g Albumax I, 3.5 μg biotin, 85 mg lactic acid, and 2.5 mg ascorbic acid.

### ZIKV stocks

Seven ZIKV strains were used. Six strains, including three African lineage strains isolated in Uganda (MR766), Nigeria (IbH 30656), and Senegal (DAK AR 41524); two Asian lineage strains isolated in Malaysia (P6-740) and Cambodia (FSS13020); and one American lineage strain isolated Puerto Rico (PRVABC59) were obtained from BEI Resources. One additional American lineage strain isolated in Brazil (HS-2015-BA-01) was graciously gifted by Dr. Mauro M. Teixeira from Albert Einstein College of Medicine. ZIKV strains were expanded in C6/36 mosquito cells in NGD 0.5X medium at 28°C. Virus-containing cell supernatant was harvested after the appearance of cytopathic effect, clarified with centrifugation, and concentrated using a Centricon Plus-70 filter (100 kDa cutoff). Conditioned media was collected from uninfected C6/36 cells grown in parallel. Aliquots were stored at −80°C.

### Plaque assay for viral titer

Viral titer was determined by plaque assay in BHK-21 cells (ATCC CCL10). Cells were infected with dilutions of viral stock or supernatant collected from infected samples and overlaid with 3.2% carboxymethylcellulose solution mixed at 1:1 with Dulbecco’s Modified Eagle Medium (DMEM) containing 4% fetal bovine serum (FBS). At 4 days post-infection (dpi), cells were fixed in ice cold methanol and stained with 0.5% crystal violet. Plaques were counted manually.

### Phylogenetic analysis and comparison of amino acid sequences

Genome sequences of the seven ZIKV strains were obtained from GenBank (NIH). Chronograms were estimated with BEAST version 2.2.129 using a general time reversible nucleotide substitution model with a discrete gamma distribution and random local clock as previously optimized for ZIKV phylogenetic analysis.

Amino acid sequences of the seven ZIKV strain RNA open reading frames were obtained from GenBank (NIH) and aligned using Clustal Omega (EMBL-EBI) (*102*). A percent identity matrix was then generated from the alignments.

### Maintenance of human embryonic stem cells (hESCs)

WIBR3 hESCs (NIHhESC-10-0079 were maintained in feeder-free conditions with StemFlex media (Gibco) on Matrigel-coated (Corning) 6-well tissue culture dishes. For passaging, cells were detached as clumps using Versene Solution (Thermo) and replated at a ratio of 1:8-1:10.

### Differentiation of hESCs to NPCs and maintenance

5 million hESCs (WIBR3, NIHhESC-10-0079) were passaged onto Matrigel-coated 6-well dishes using Accutase (Thermo) directly in NGD 0.5X media containing dorsomorphin (2.5 μM, Stemgent), basic fibroblast growth factor (bFGF) (10 ng/mL, Thermo), insulin (1:500), and Y-27632 Rho-associated protein kinase (ROCK) inhibitor (10 mM, Stemgent). Media was replaced daily without addition of ROCK inhibitor. After one week, bFGF was removed and the media was replaced every day with fresh NGD 0.5X media with dorsomorphin and insulin for an additional 7-10 days. When rosette lawns were observed throughout the culture (approximately day 15), cells were passaged 1:1 using Accutase. ROCK inhibitor was added to the medium during each of the first 3 passages. Initial passaging was at no more than a 1:2 ratio, followed by 1:3 to 1:6 every 5 days. After the initial passage, neural progenitors were expanded and maintained in NGD 0.5X with bFGF and insulin.

### 2D cell culture ZIKV infection and sample preparation

Infection of 2D cell culture was carried out at an MOI of 1 based on the number of cells seeded the day prior. Inoculum was prepared by diluting viral stock in cell culture media, with the volume differences among viral stock compensated for by conditioned media. Mock inoculum contained conditioned media equal to the highest volume of viral stock added. Infections were allowed to occur for 1 hr at 37°C, after which the inoculum was replaced with fresh cell culture media. Samples were collected at 48 and 72 hours post-infection (hpi).

For biochemical assays, cells were detached with Accutase and centrifuged to form a pellet. After removing the supernatant, cell pellets were snap frozen in liquid nitrogen.

### Cerebral organoid culture

Cerebral organoids were generated from hESCs based on a published protocol by Lancaster *et al*. (*103*), and cultured with the STEMdiff Cerebral Organoid Kit (Stem Cell Technologies). Briefly, hESCs (WIBR3, NIH ESC-10-0079) were placed in mTeSR1 media (Stem Cell Technologies) with 10 μM ROCK inhibitor (Thermo) one day prior to seeding. Embryoid bodies (EBs) were generated by seeding 9,000 hESCs in a 96-well U-bottom ultra-low attachment plate with EB formation media and 10 μM Y-27632 ROCK inhibitor. Additional EB formation media was added to the wells on days 2 and 4. EBs were then transferred on day 5 into 24-well ultra-low attachment plates with Neuroinduction media. On day 7, EBs were embedded in 15 μl Matrigel droplets using sheets of dimpled parafilm and incubated for at least 30 min at 37°C. EBs were then transferred into 6 cm dishes with 4 ml of Expansion media. On day 10, the organoids were transferred into 6-well ultra-low attachment plates with 3 ml Maturation media and cultured on an orbital shaker set to 65 rpm. Maturation media was replaced every 3-4 days.

### Cerebral organoid ZIKV infection and sample preparation

Cerebral organoids were infected at 21 days post-EB formation at 500 focus-forming units/200 μl media per organoid, or an MOI of 0.002 based on the estimated number of cells on the surface of an organoid. Inoculum was prepared by diluting viral stock in Maturation media, with the volume differences among viral stock compensated for by conditioned media. Mock inoculum contained conditioned media equal to the highest volume of viral stock added. Infections were allowed to occur for 24 hrs at 37°C on an orbital shaker set to 65 rpm, after which the inoculum was replaced with fresh Maturation media.

Cerebral organoids were collected at 7, 14, and 21 dpi. Twenty-four hours prior to a collection time point, the media was replaced with 333 μl of Maturation media per organoid so that the collected supernatant would reflect 24 hrs of virus production. For biochemical assays, organoids were washed with PBS and the Matrigel was removed as much as possible before snap freezing with liquid nitrogen. Samples were stored at −80°C until needed and lysates were created by incubating samples on ice for 10 min with cell lysis buffer (Cell Signaling Technology).

Organoid lysates were then separated for protein and RNA assays. For immunofluorescence studies, organoids were fixed in 4% PFA in PBS for 30-45 min at room temperature. After fixation, organoids were washed three times with PBS.

### Cerebral organoid cryosectioning

After fixation, cerebral organoids were cryoprotected with sequential incubations in 15% and 30% w/v sucrose (Sigma) in PBS overnight at 4°C. Organoids were then embedded in OCT compound (Fisher) and snap frozen in 2-methyl-butane (Sigma) cooled in liquid nitrogen. Organoid blocks were stored at −80°C and cryosectioned at 20 μm with a Leica CM1850 Cryostat. Sections were captured onto Superfrost Plus microscope slides (Fisher) and allowed to fully dry at room temperature before being stored at −80°C.

### Immunofluorescence staining

Organoid sections were washed in PBS prior to staining, then permeabilized and blocked with 0.3% Triton X-100 in 3% normal goat serum in PBS for 1 hr at room temperature. The sections were stained for the following antibodies: rabbit monoclonal anti-SOX2 (1:500, Cell Signaling Technology), chicken polyclonal anti-MAP2A/B (1:10,000, EnCor Biotechnology), and mouse monoclonal J2 anti-double stranded RNA (1:2000, SCICONS) in blocking buffer overnight at 4°C. After primary antibody incubation, sections were washed three times with PBS and 0.25% Triton-X (v/v) in PBS (PBST). For secondary antibody staining, goat anti-mouse, anti-chicken, or anti-rabbit antibodies were conjugated with Alexa Fluor Dyes (1:1000, Invitrogen) and DAPI (1 μg/ml, Invitrogen) in blocking buffer. Incubation time was 1 hr at room temperature. After secondary antibody incubation, sections were washed three times with PBST, and then three times with PBS. #1.5 thickness coverslips were mounted with ProLong Diamond antifade mountant (Invitrogen), and the samples were allowed to cure overnight at room temperature before being stored at 4°C.

### Fluro-Jade C (FJC) staining

The Fluoro-Jade C Staining Kit (Biosensis) was used according to the manufacturer’s protocol to stain for degenerating neurons (*52*). Organoid sections were washed in PBS prior to staining. #1.5 thickness coverslips were mounted with Cytoseal XYL (Thermo) after staining, and the samples were allowed to cure overnight before being stored at room temperature.

### MitoSOX Red staining

Vero cells (ATCC CCL-81) were infected with ZIKV at an MOI of 1 as described above. At 24 hpi, MitoSOX Red staining was performed per the manufacturer’s protocol.

### Trolox treatment

Trolox (Sigma) was reconstituted in DMSO (Sigma) before being diluted to a final concentration of 100 μM in Maturation media. Control media contained DMSO equal to the same volume of Trolox added. After cerebral organoid ZIKV infection as described above, the inoculum was replaced with either the treatment or control media. Organoid collection and preparation then occurred as described above.

### Microscopy and image analysis

Darkfield images were acquired with a Leica DFC310 FX camera. Organoids were manipulated by swirling and tilting the plate to ensure that they were not compressed against each other or the walls of the well before image acquisition. Confocal microscopy was performed using an Olympus FV1200 laser scanning confocal microscope. Images were acquired using both a 30X and 60X oil-immersion objective.

Microscopy images were analyzed using Fiji (*104*). Organoid size was determined by measuring the area of nine organoids per condition in darkfield images. Each organoid was individually tracked over time. For confocal images, each quantification was done on at least three independent fields of view per condition. The StarDist 2D plug-in (*105*) was used to help automatically detect and segment tightly packed nuclei in organoid images. Incorrectly segmented nuclei and ones touching the edge of the image were removed before analysis. For NAF and percentage SOX2+/J2+ analysis, a threshold was set so that only nuclei with SOX2 staining above the mean intensity of all measured nuclei in each field of view were quantified. At least 500 nuclei were quantified for each condition. MAP2 area was measured on binarized images with the MAP2 threshold normalized to Mock organoid images as previously published (*106*). This value was then divided by the total number of nuclei in the same image, excluding any SOX2+ nuclei. Each field of view contained at least 250 nuclei.

### Immunoblotting

Snap frozen cerebral organoids and cell pellets were lysed by dispersing in 1X RIPA buffer (50 μl/organoid or 10^6^ cells, Thermo-Fisher Scientific) with protease inhibitors for 15 min on ice with occasional mixing. Lysates were centrifuged at 14,000×g for 10 min at 4°C. The supernatants were collected and heated at 65℃ in SDS-PAGE sample buffer for 15 min prior to SDS-PAGE in 4-15% or 4-20% polyacrylamide gels. Separated proteins were then transferred onto polyvinylidene difluoride (PVDF) membranes. The blots were first stained with Direct Blue 71 (DB71) (Sigma) so that total protein could be used as the loading control. Total protein staining has been shown to be equivalent or superior to housekeeping protein-based methods (*107*). After removing the DB71 stain, the blots were blocked with 5% non-fat milk in Tris-buffered saline with 0.1% Tween 20 (TBST) (Gibco). The blots were then washed three times with TBST for 10 min at room temperature, and incubated with primary antibodies in TBST with 1.5% bovine serum albumin (BSA) overnight at 4°C. The primary antibodies used in this study were as follows: rabbit anti-NOXA (1:1000, Cell Signaling Technology), rabbit anti-cleaved caspase-3 (1:1000, Cell Signaling Technology), mouse anti-Bcl-2 (1:1000, Cell Signaling Technology), rabbit anti-eIF2αP (1:1000, Abcam), rabbit anti-ATF4 (1:1000, Cell Signaling Technology), goat anti-GLRX1 (1:400, R&D Technologies), rabbit anti-Zika virus envelope protein (1:3000, GTX), rabbit anti-IFIT1 (1:1000, Cell Signaling Technologies), mouse anti-STAT2 (1:100, Santa Cruz Biotechnology), and rabbit anti-ATF3 (1:1000, Cell Signaling Technologies).

After primary antibody staining, the blots were washed three times with TBST, and then incubated with ddsHRP-conjugated secondary antibodies (1:7500, Promega) in TBST with 1.5% BSA for 1 hr at room temperature. Three more TBST washes were done following secondary antibody staining, before the blots were developed with ECL reagents and imaged in a Syngene G:BOX. Western blot images were analyzed using Fiji (*104*).

### Bulk RNA isolation and sequencing

Total RNA was extracted from organoid lysates using the RNeasy Mini Kit (QIAGEN) following the manufacturer’s instructions. Samples were quantified with a NanoDrop (Thermo) before storing at −80°C. Samples were submitted to the MIT Genomics and Bioinformatics core for RNA purification, library preparation, and sequencing. RNA-seq data were used to quantify transcripts from the hg38 human assembly with the gencode version 46 basic annotation plus virus genomes (ZVMA:KX377336.1; ZVPR: KX087101.3; ZVUG: U963573.2), using the nf-core/maseq workflow revision 3.14.0 (*108*). Gene level summaries were prepared from the star_salmon quantitation using tximport version 1.44.0 (*109*) running under R version 4.4.1 (R Core Team 2021 with tidyverse version 2.0.0 (*110*). Differential expression analysis was done with DESeq version 1.44. Gene set enrichment analysis (GSEA) and gene ontology analysis were then performed by the Bioinformatics & Computing Core Facility of the Swanson Biotechnology Center, Koch Institute, at the Massachusetts Institute of Technology, Cambridge MA.

### Single cell RNA sequencing

Mock, ZIKV-PR, ZIKV-MAL, and ZIKV-UG infected organoids at 7 dpi were manually dissociated into a single cell suspension using the Embryoid Body Dissociation Kit (Miltenyi Biotec). The manufacturer’s protocol was followed with these adjustments: 1 ml of enzyme mix was used per 3 organoids, 1 ml of ice-cold Maturation media was used to wash the pre-separation filter, and the cell suspension was centrifuged at 300×g for 10 min to better pellet neurons.

scRNA-seq was performed using the Chromium Next GEM Single Cell 3’ Kit v3.1 (10x Genomics) following the manufacturer’s protocol. Single cell suspensions were loaded onto the Chromium Controller (10x Genomics) for droplet formation. scRNA-seq libraries were prepared using the Chromium Single Cell 3’ Reagent Kit (10x Genomics). Samples were sequenced by the MIT Genomics and Informatics Facilities core. Raw reads were then mapped to a human reference genome including the ZIKV genomic sequences listed as individual genes by the same core facility. Mapped data were analyzed using Seurat.

### Single cell RNA-seq analysis

#### Data Acquisition and Preprocessing

The CellRanger Single-Cell Software Suite (cellranger-6.0.1) (https://www.10xgenomics.com/support/software/cell-ranger/latest/getting-started/cr-what-is-cell-ranger) was used to perform sample demultiplexing, barcode processing and single-cell 3′ gene counting. Complementary DNA reads were aligned to the GRCh38 reference genome, which was slightly modified to include three reference alignments for the ZIKA viruses (vir-ZVPR, vir-ZVUG, and vir-ZVMAL) at the end. CellRanger outputs from individual samples were obtained and each dataset was read using the Read10X function in R to load the gene-barcode matrices.

#### Seurat Object Initialization

Four Seurat objects were initialized using the raw data, setting parameters for minimum cells (min.cells = 3) and minimum features (min.features = 200)(*111*). Each Seurat object was labeled with its respective experimental condition: “Mock”, “Puerto Rico”, “Malaysia”, and “Uganda”.

#### Data Integration and Normalization

The Seurat objects were merged into a single integrated dataset using the merge function, creating a unified project. The merged dataset underwent normalization (normalization.method = “LogNormalize”, scale.factor = 10000), feature selection (FindVariableFeatures, selection.method = “vst”, nfeatures = 2000), and data scaling (ScaleData). Principal Component Analysis (PCA) was conducted to reduce dimensionality, and uniform manifold approximation and projection (UMAP) was performed for visualization (RunUMAP) using 30 dimensions based on Elbow plot result. Subsequently, the dataset was clustered using FindNeighbors and FindClusters functions.

#### Quality Control and Filtering

Quality control metrics such as the percentage of mitochondrial genes (percent.mt) and log-transformed gene counts per cell (log10GenesPerUMI) were computed and visualized using violin plots (VlnPlot). Cells were filtered based on criteria of percent.mt <= 25 and log10GenesPerUMI > 0.75 to remove low-quality cells and retain high-quality data for downstream analysis.

#### Integration and Visualization

Data integration across conditions was performed using SCT normalization (SCTransform, vars.to.regress = c(“percent.mt”)), feature selection for integration (SelectIntegrationFeatures, nfeatures = 3000), and identification of integration anchors (FindIntegrationAnchors). Integrated data visualization included PCA and UMAP plots to explore global and local similarities among conditions.

#### Viral RNA Expression Analysis

To analyze viral gene expression within the Seurat object, violin plots were generated to visualize the expression levels of specific viral genes across different conditions. Three individual violin plots were created for the viral genes vir-ZVMAL, vir-ZVPR, and vir-ZVUG, grouped by the Condition metadata (“Mock”, “Malaysia”, “Puerto Rico”, “Uganda”).

#### Cell Type Assignment of Clusters

The integrated Seurat object was utilized for cell type assignment. Initially, the default assay was set to “RNA” using DefaultAssay(seurat.integrated) <-“RNA”. Clusters were identified based on the integrated nearest neighbor graph resolution 0.4 by assigning cluster identities through Idents(object = seurat.integrated) <-‘integrated_snn_res.0.4’.

Subsequently, marker genes for each cluster were determined using the FindAllMarkers function. This analysis was restricted to positive markers with a minimum detection percentage of 25% (min.pct=0.25) using the parameters only.pos=TRUE and min.pct=0.25. The resulting stronger signature marker genes were validated by violin and feature plots and then compared to the literature for celltypes identification. Additionally, a bubble plot was created to visualize the expression levels of selected marker genes across different conditions using the DotPlot function. Neuronal progenitor cells (NPCs) were specifically identified by the expression of SOX2, a well-established biomarker for NPCs(*112–114*). Intermediate progenitor cells (IPCs) were identified by the expression of TOP2A, CCNB2, and CENPF, which are associated with cell cycle progression and progenitor states(*114*). Choroid plexus cells were identified by the expression of TTR, PCP4, and HTR2C(*115–117*). Radial glial cells, critical for neurogenesis and acting as scaffolds for migrating neurons, were identified by the expression of PTN (*112, 118*). Mature neurons were identified by the expression of MAP2 and STMN2, markers associated with neuronal structure and function (*114, 119, 120*). Midbrain-Hindbrain cells were identified by the expression of NEFL and MEIS2, reflecting their role in neuronal maturation (*121–123*). Astrocytes were assigned by the expression of S100A10 (*124*).

To visualize the distribution of cells across various cell types under different experimental conditions, we categorized the experimental conditions in the Seurat-integrated object and concatenated the cluster identities with these conditions to form unique identifiers for each cluster-condition combination. These identifiers were set as the active identities in Seurat, allowing us to generate a frequency table of the combined identifiers. This table was subsequently converted into a dataframe, and the combined identifiers were split into separate columns for clusters and conditions using the tidyr::separate function. The resulting data frame was used to create a bar plot with ggplot2. In this plot, the x-axis represents the different clusters, the y-axis indicates the number of cells, and the fill colors differentiate the experimental conditions.

#### Differential Expression Analysis

Differential expression analysis was performed to identify genes that were differentially expressed across conditions. This analysis was conducted separately for “Puerto Rico” vs. “Mock”, “Malaysia” vs. “Mock”, and “Uganda” vs. “Mock” comparisons within each celltype using the FindMarkers function (logfc.threshold = 0).

#### Gene Set Enrichment Analysis (GSEA)

Ranked gene lists (rnk files) were generated from differential expression results using avg_log2FC values. The process involved iterating through each cluster of differential expression results for the conditions MAL, PR, and UG, extracting the avg_log2FC values, and renaming the columns to reflect the specific condition and cluster (e.g., Ug_v_mock.Neuronal_progenitor_cells.rnk, PR_v_mock.Neurons.rnk, etc.). These avg_log2FC values were stored in individual data frames for each cell type, and the data frames were subsequently merged into a single data frame for each condition.

For each merged data frame, the gene name and corresponding avg_log2FC values were selected and saved as rnk files, ensuring only complete cases were included. These rnk files were then used in pre-ranked Gene Set Enrichment Analysis using javaGSEA version 4.2.3 with msigDb version v20203.2 and Gene Ontology Biological Process (c5bp). After running the GSEA, the resulting data files were parsed into tabular formats with custom scripts.

### Statistical analyses

All statistics were performed using GraphPad Prism 9.0.2 software. Statistical significance was determined using two-way ANOVA with Tukey’s multiple comparisons test.

## List of Supplementary Materials

**Figure S1.** Schematic diagram of an organoid ventricle, showing cell types. RG: radial glia; NPC: neuronal progenitor cells; IPC: intermediate progenitor cells.

**Table S1.** Cell counts of individual cell types identified for single cell RNA-seq analysis.

## Author contributions

ATH: experimental design, performed experiments, writing, RNA-seq analysis, data interpretation, figure preparation

YZ: experimental design, performed experiments, image acquisition and analysis, writing, data interpretation, figure preparation

J-JC: experimental design, data interpretation, protein immunoblotting experiments, expertise in unfolded protein response and integrated stress response, manuscript editing

JMA: organoid preparations, experimental design, performed experiments, data interpretation

AR: stem cell maintenance, stem cell and NPC preparations, experimental design, data interpretation

VL: technical assistance with organoid preparations, experimental design, performed experiments

CAW: bioinformatic analysis and design, data interpretation, figure preparation, manuscript review

YSV: bioinformatic analysis and design, data interpretation, figure preparation, manuscript review

DA: single cell RNA-seq analysis, figure preparation, data interpretation

TL: preparation and maintenance of stem cells and NPC

RJ: project design, funding, supervision, manuscript review

LG: project design, funding, supervision, data analysis, writing

