## Supplemental Figure 1 for "Oxidative Stress and Interferon Signaling Drive Differential Pathogenesis of Ancestral and Contemporary Zika Viruses in Human Cerebral Organoids"

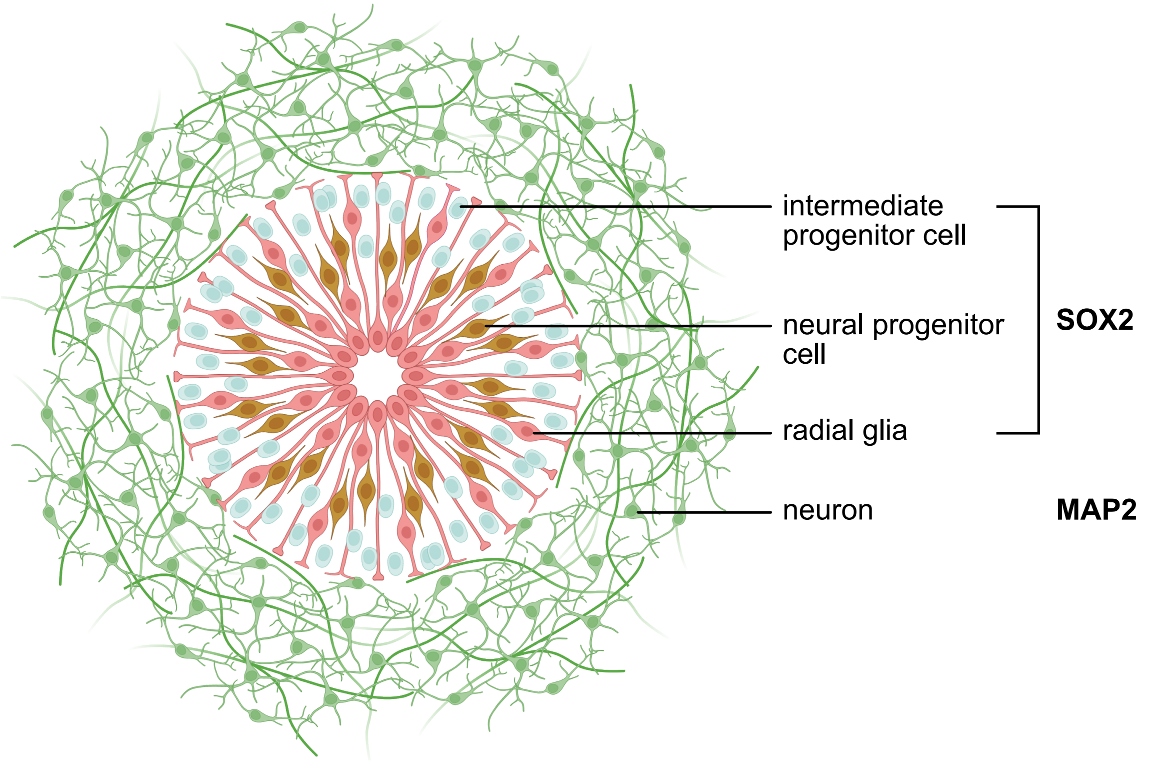


**Figure S1.** Schematic representation of an organoid ventricle. Radial glia, neural progenitor cells, and intermediate progenitor cells stain with anti-SOX2 antibodies, while neurons stain with anti-MAP2 antibodies. This image was created in BioRender.
