## Supplemental Table 1 for "Oxidative Stress and Interferon Signaling Drive Differential Pathogenesis of Ancestral and Contemporary Zika Viruses in Human Cerebral Organoids"

| | NPC | Neurons | IPC | Radial Glia | MB-HB | Chor Plex | Astro | ISG-High | NPC-Like |
|---|---|---|---|---|---|---|---|---|---|
| Puerto Rico | 1281 | 670 | 372 | 332 | 388 | 128 | 84 | 198 | 129 |
| Uganda | 3170 | 745 | 704 | 190 | 161 | 145 | 57 | 413 | 155 |

**Table S1.** Cell counts of individual cell types identified for single cell RNA-seq analysis.
